# Psilocybin Promotes Cell-Type-Specific Changes in the Orbitofrontal Cortex Revealed by Single-Nucleus RNA-seq

**DOI:** 10.1101/2024.01.07.573163

**Authors:** Ziran Huang, Xiaoyan Wei, Yihui Wang, Jing Tian, Jihui Dong, Bo Liang, Lin Lu, Wen Zhang

## Abstract

Recent clinical breakthroughs hold great promise for the application of psilocybin in the treatments of psychological disorders, such as depression, addiction, and obsessive-compulsive disorder. Psilocybin is a psychedelic whose metabolite, psilocin, is a 5-HT_2A_ receptor agonist. Nevertheless, the underlying mechanisms for the effects of psilocybin on the brain are not fully illustrated, and cell type-specific and circuit effects of psilocybin are not fully understood. Here, we combined single-nucleus RNA-seq with functional assays to study the long-term effects of psilocybin on the orbitofrontal cortex (OFC), a brain region vulnerable to psychological disorders such as depression. We showed that a single dose of psilocybin induced long-term genetic and functional changes in neurons of the OFC, and excitatory and inhibitory neurons collectively reduced circuit activity of the brain region. Knockdown of 5-HT_2A_ receptor in deep layer excitatory neurons abated psilocybin-induced functional changes and the anti-depressant effect. Together, these results showed the cell type-specific mechanisms of psilocybin and shed light on the brain region difference in the effect of psychedelics.

## Introduction

The increase in the prevalence of depression globally has put a great burden on the global economy ^1^. Compared with the last decade, the prevalence of depression disorders increased by more than 120 million globally ^1^, and the COVID-19 pandemic further exacerbated such a trend^2^. On the other hand, treatments with traditional anti-depressants were still unsatisfactory ^3, 4^. Such a gap urges the development of novel medications with higher efficiency. Recently, psychedelics have been proposed as a fast-acting anti-depressant, and clinical trials suggest their promising roles in treating addiction, obsessive-compulsive disorder (OCD), and post-traumatic stress disorder (PTSD) ^5–11^. However, similar to traditional anti-depressants, the underlying cellular and circuit mechanisms of psychedelics are still not fully understood. Considering the hallucinogenic effects of psychedelics, the lack of mechanistic understanding could limit their clinical use, since they might require extra costs for medical supervision and safety monitoring during treatments.

Studies have shown that psychedelics, such as psilocybin, lysergic acid diethylamide (LSD), and N,N-Dimethyltryptamine (DMT), activate the 5-HT_2A_ receptor, which activates G_αq_ and β-arrestin signaling pathways and then lead to downstream effects ^12^. Evidence also suggested that psychedelics, such as psilocybin, induced plasticity changes in neurons, such as an increase of spines of excitatory neurons ^13–15^. Meanwhile, brain functions require concerted activities of both the excitatory and inhibitory neurons, which not only underlies the plasticity of the brain, but the shift of such balance between excitation and inhibition also could predispose neurological disorders, such as autism and schizophrenia ^16, 17^. In the cortex, inhibition is mostly mediated by GABAergic interneurons, which exhibit both genetical and physiological diversity ^18^. Such diversity requires cell type-specific investigation of interneuron functions in both physiological and pathological conditions, as both brain region and layer-specific effects of interneurons on the output exist in the cortex ^19, 20^. However, the effects and mechanisms of psychedelics action on different types of cortical neurons are not fully illustrated, which further dampens our understanding of the mechanisms of effects of psychedelics on neuron circuits in the brain.

Furthermore, while reports suggest that psychedelics increase the dendritic spines in the frontal cortex and somatosensory cortex ^21, 22^, studies also found that such effect might not be consistent across brain regions. For example, the default mode network (DMN) is a conserved brain network vulnerable to brain disorders, which showed hyperconnectivity in depressive patients ^23^. Single-dose application of psychedelics to healthy subjects and patients with treatment-resistant depression decreased DMN activity and functional connectivity ^24, 25^. The orbitofrontal cortex (OFC) is a brain region that showed extensive enervation to brain regions in the DMN in both human and rodents ^26, 27^, and psilocybin and other anti-depressants reduced the activity of the OFC or the functional connectivity of the OFC with brain region in the DMN ^28, 29^; however, the underlying mechanisms of such functional changes is unclear.

To illustrate the impact of psychedelics on the brain and its possible cell type-specific effect, in the present study, we combined single-nucleus RNA-seq, electrophysiology, and behavior tests to study the effect of single-dose application of psilocybin on gene expression and the function of cortical neurons in the OFC. Additionally, we also explored their possible contributions to the anti-depressant effect of psilocybin.

## Results

### Single-cell landscape of the orbitofrontal cortex after psilocybin injection

To understand how psilocybin induced long-term changes in the cortex, we injected a single dose of psilocybin (1 mg/kg, intraperitoneally [*i.p.*]) into adult male mice (8 -12 wks old) with mice injected with vehicle (saline) as control. Studies have shown that such dosage induced a sharp rise of head-twitch response, which is considered as hallucinatory behavior in rodents, dwindled to normal ∼ 3 hrs after injection in mice, while cortical neurons still showed persistent functional changes afterward ^30^. We did not observe a difference in the head-twitch responses between saline- and psilocybin-injected (Saline and Psilocybin, respectively) mice 24 hours after injection (**Figure S1A** and **B**), and these mice showed no difference in the open field test (**Figure S1C**). We dissected the brain region of the orbitofrontal cortex (OFC) 24 hrs after injection, then extracted the nucleus of cells to perform single-nucleus RNA sequencing (snRNA-seq, **Figure 1A**). We generated data on 25237 cells post-quality control with 12246 cells from Saline mice and 12991 cells from Psilocybin mice (N = 4 mice per group, **Table S1**). We identified 20 clusters of cells, all of which contain cells from both Saline and Psilocybin mice (**Figure 1B, Figure S2A,** and **Table S1**). These clusters were grouped into 3 major cell types based on known cortical cell markers ^31–34^, including Glutamatergic, GABAergic, and non-neuronal cells (**Figure 1C** and **Figure S2B**).

**Figure 1.**
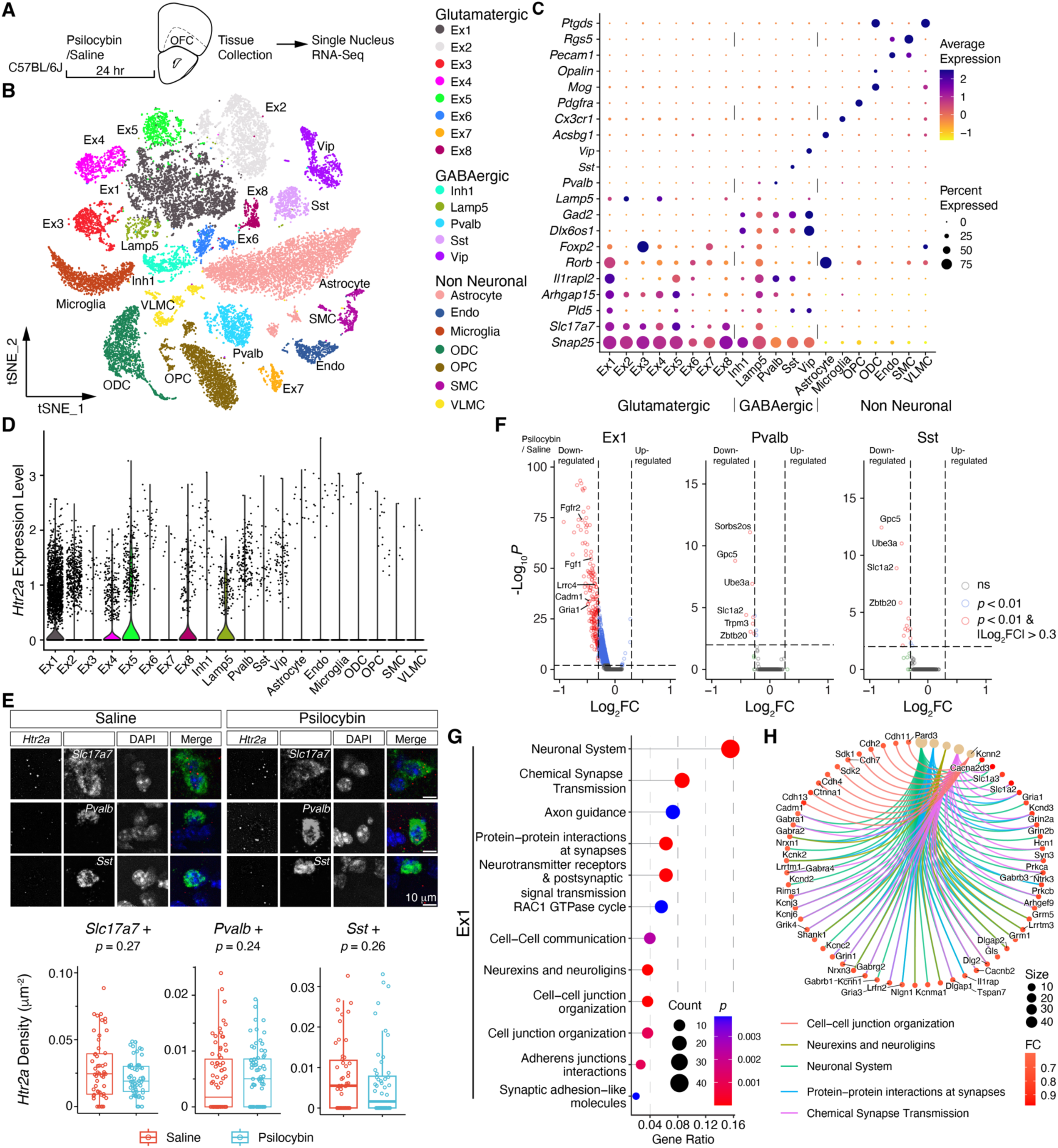
Single nucleus RNA sequencing of cells in the OFC of saline- and psilocybin-injected mice. **(A)** The sampled for single nucleus RNA sequencing (snRNA-seq) experiment was performed 24 hrs after *i.p.* injection of psilocybin (Psilocybin) or the vehicle (Saline). **(B)** Visualization of cell clusters in the OFC with t-distributed stochastic neighbour embedding (*t*-SNE). Dots represent individual cells, which were clustered into 3 major groups, 20 subgroups (Saline, n = 12246; Psilocybin, n = 12991; N = 4/group). **(C)** Dot plot of the expression of cell-type markers in clusters. **(D)** Violin plot of *Htr2a* expression in clusters. **(E)** RNAScope *in situ* hybridization analysis of *Htr2a* in layer 5 excitatory (*Slc17a7*+), inhibitory PV+ (*Pvalb*+), and inhibitory SST+ (*Sst*+) neurons of the OFC in Saline and Psilocybin mice. Top panels, characteristic images of *Htr2a* mRNA puncta in *Slc17a7*+, *Pvalb*+, and *Sst*+ cells in layer 5 of OFC. Bottom boxplots, the comparisons of the density of *Htr2a* puncta in the three cell types (*p*, Wilcoxon test; *Slc17a7*+: Saline, n =52; Psilocybin, n = 55; *Pvalb*+: Saline, n = 74; Psilocybin, n = 68; *Sst*+: Saline, n = 50; Psilocybin, n = 52; N = 3/group). **(F)** Volcano plots of the differential gene expression analysis of the Ex1, Pvalb, and Sst cluster of snRNA-seq data between Psilocybin and Saline mice. **(G)** Reactome pathway-based gene set enrichment analysis of differential gene expressions in the Ex1 cluster between Psilocybin and Saline mice. **(H)** Network analysis of genes showing differential expression in Psilocybin mice with top annotated Reactome pathway categories of the Ex1 cluster.

Psilocybin is metabolized and dephosphorylated to psilocin in the body, which is an agonist of the 5-hydroxytryptamine receptor subunit 2A (5-HT_2A_R). In the medial frontal cortices of primates and rodents, 5-HT_2A_Rs are densely expressed in layer 5 pyramidal neurons and interneurons ^35, 36^. We examined the expression of the mRNA (*Htr2a*) of 5-HT_2A_R in cell clusters and 5-HT_2A_R in cells of the OFC. We found both *Htr2a* and 5-HT_2A_R majorly expressed in the Glutamatergic and GABAergic neurons but not in those non-neuronal cells of the cortex (**Figure 1D and Figure S3**). Of the Glutamatergic cell clusters, the Ex1 cluster also showed strong expression of *Pld5*, which showed high abundance in the deep layers of the OFC (**Figure 1C** and **Figure S2B**) ^37^. We then quantified the expression of the mRNAs of *Htr2a* in the OFCs with RNAscope *in situ* hybridization (**Figure 1E** and **Figure S4**) in layer 5 *Slc17a7*-, *Pvalb*-, and *Sst*- positive cells in the OFC. We found no difference in *Htr2a* expression between Saline and Psilocybin mice, which suggests that psilocybin might not change *Htr2a* transcription in the OFC.

### Psilocybin induces cell-type-specific gene expression changes in the OFC

To characterize the effect of psilocybin on gene expressions in the OFC, we performed differential gene expression analyses of cell clusters in the OFC. We found that the cell cluster Ex1, the purported layer 5 pyramidal neurons, showed significant changes in gene expression (**Figure 1F**). Meanwhile, both Pvalb and Sst clusters showed less differential expression of genes than the Ex1 cell cluster (**Figure 1F**). These results suggest psilocybin-induced cell-type-dependent gene expression change. For the Ex1 cluster, we performed gene enrichment analysis based on the Reactome pathway database ^38^. We found the majority of genes showing differential expression in Psilocybin mice enriched in synaptic transmissions and cell-cell interactions (**Figure 1G** and **Table S2**), such as *Gria1* (**Figure 1H** and **Figure S5**), suggesting changes in synapse functions.

### Psilocybin reduced cell-cell interactions among neurons in the OFC

The gene enrichment analysis showed gene expression changes of multiple signaling pathways involved in cell-cell interactions (**Figure 1G**), especially in the synapse formation and maintenance of the Ex1 cluster. To identify the major signaling changes in these intercellular communications between cell clusters, we utilized CellChat ^39^ to analyze snRNA-seq results to characterize and compare the inferred cell-cell communication networks between clusters of Saline and Psilocybin mice.

We calculated the relative interaction strength of both outgoing and incoming interactions of clusters and found that the majority of clusters showed reductions in those interactions of Psilocybin mice (**Figure 2A**). We also ranked the significant signaling pathways based on the differences in the overall information flow within the inferred networks of clusters between Saline and Psilocybin mice (**Figure 2B**). We compared the changes in cell-cell interaction signaling pathways between each cluster, and found that for the majority of cell-cell interaction between clusters, Psilocybin mice showed reductions in both the numbers of interactions between a given cluster-pair and the calculated interaction strength between such pair (**Figure 2C**).

**Figure 2.**
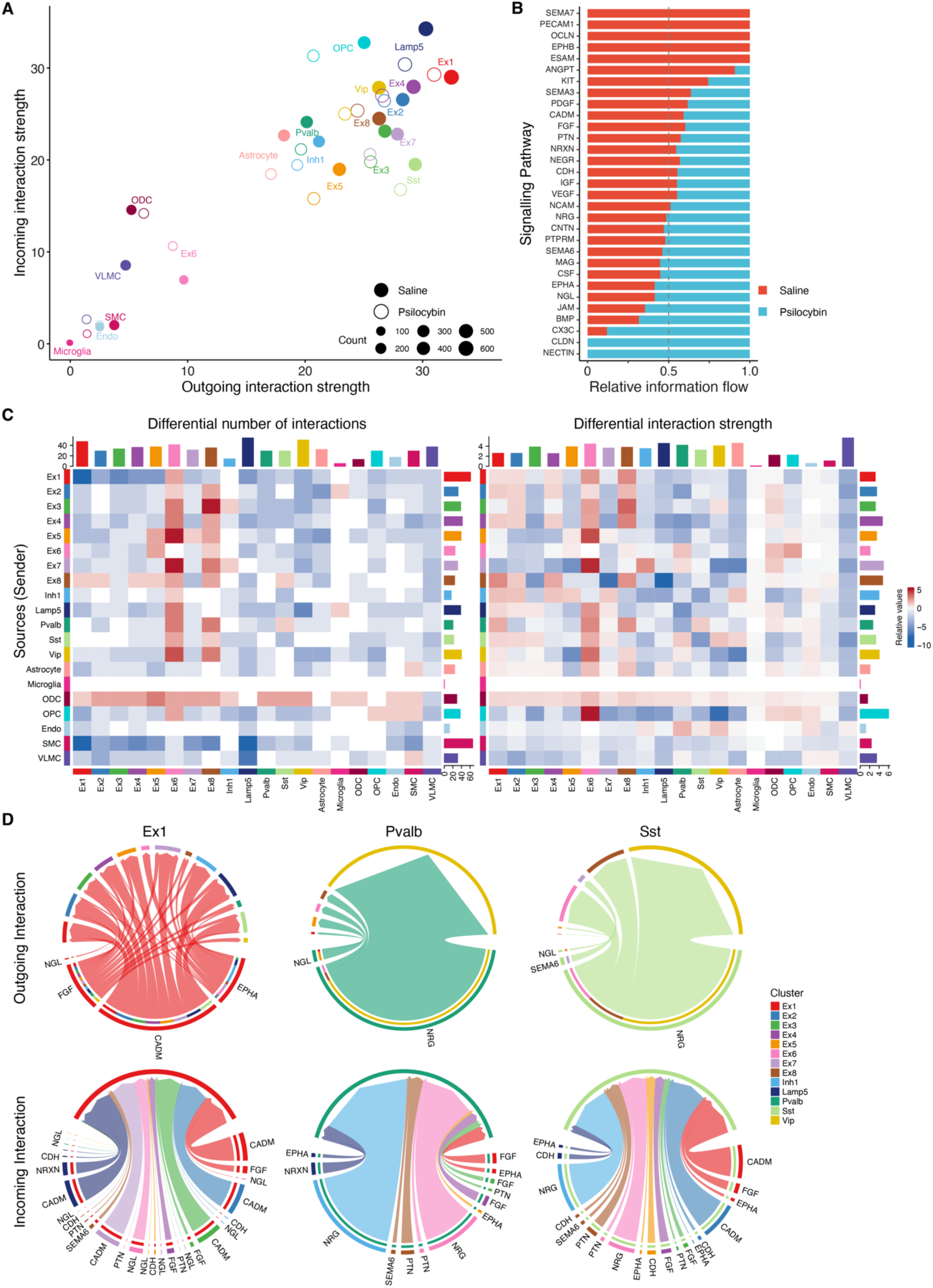
Psilocybin affected cell-cell interactions in the OFC. **(A)** Comparison of the outgoing and incoming cell-cell interaction strength of clusters between Saline and Psilocybin mice with CellChat and snRNA-seq. **(B)** Stacked plot of the major signaling pathways different between Saline and Psilocybin mice based on the overall information flow of cell-cell interaction signaling pathways between these mice. The top signaling pathways colored red are enriched in Saline mice, and these colored green were enriched in Psilocybin mice. **(C)** Heatmaps of the changes in the number of interactions (left panel) and the interaction strength (right panel) between clusters of Psilocybin compared with Saline mice. For heatmaps, the top bar plot represents the sum of column of values displayed in the heatmap (incoming signaling). The right bar plot represents the sum of row of values (outgoing signaling). **(D)** Chord diagrams of the changed cell-cell interaction signaling pathways of Ex1, Pvalb, and Sst clusters with other neuronal cell clusters in Psilocybin mice. Arrows in chord diagrams indicate the target clusters of cell-cell interactions. For the parts of the chord diagrams with double arcs, the inner ones indicate the targets that receive signal from the corresponding outer arcs, and the arc size is proportional to the signal strength received by the target.

We further analyzed all the interactions sending or receiving from the Ex1, Pvalb, and Sst clusters to other neuron clusters. We found that the affected cell-cell interaction pathways accumulated into 9 cell-cell interaction signaling pathways (**Figure 2D and Figure S6**), namely CADM, CDH, EPHA, FGF, NGL, NRG, NRXN, PTN, SEMA6 signaling pathways (**Figure S7** - **15**). Consistent with the change of global cell-cell interaction in clusters (**Figure 2C**), the Ex1, Pvalb, and Sst clusters showed more down-regulated pathways in Psilocybin mice than up-regulated (**Figure S6**). These down-regulated pathways in Psilocybin mice (**Figure 2D** and **Figure S5**) were important for synapse functions, especially excitatory synapse formation and maintenance ^40–48^. Combined with the gene set enrichment analysis of differential gene expression of the Ex1 cluster (**Figure 1G** and **H**), these analyses strongly suggest psilocybin induced changes in neuronal activity and synaptic transmissions of the Ex1 cluster.

### Psilocybin induced a reduction of activity in the OFC

To understand the functional changes of the OFC after psilocybin injection, we characterized the electrophysiological properties of neurons in the OFC. We recorded synaptic and intrinsic properties of the layer 5 Glutamatergic pyramidal neurons, GABAergic PV+, and GABAergic SST+ neurons in the OFC. Consistent with the differential gene expression analyses of cell clusters, Psilocybin mice showed cell type-specific changes in activities. The layer 5 pyramidal neurons showed reduced the output as less firing was observed (**Figure 3A**), while we did not observe a change in membrane properties or the properties of action potentials (**Figure S16A**). We also recorded spontaneous EPSCs, and consistent with a reduction in *Gria1* expression in the Ex1 cluster (**Figure S5A**), we observed decreased excitatory transmission (**Figure 3B**). On the other hand, the spontaneous IPSCs of pyramidal neurons did not change (**Figure S17A**). These results showed a decrease in synaptic activities and the output of pyramidal neurons.

**Figure 3.**
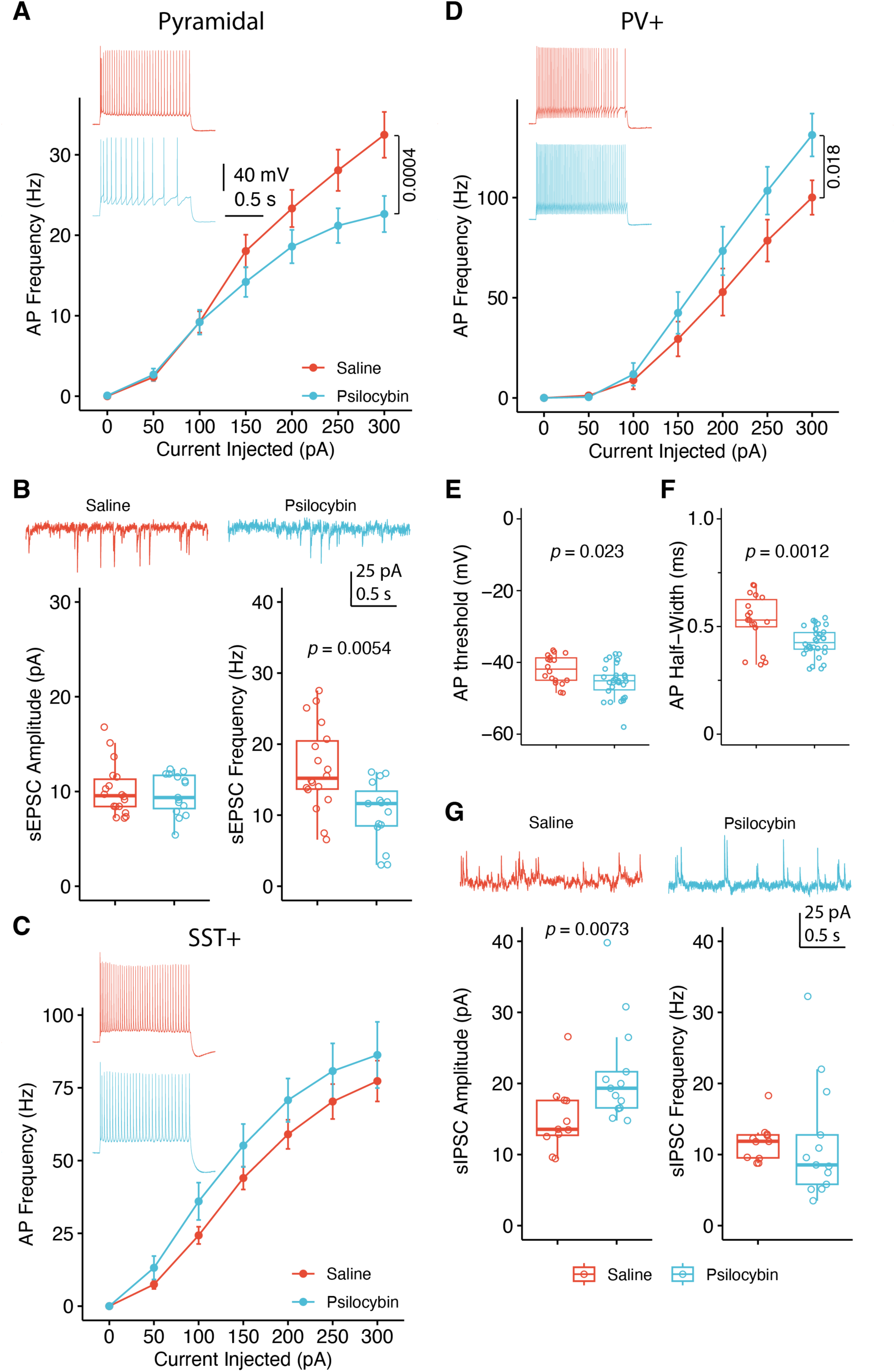
Psilocybin induced cell-type specific changes of neuronal activity in the OFC. **(A)** F-I plot shows reduced excitability of excitatory layer 5 pyramidal neurons in psilocybin mice (Two-way ANOVA, F_(3, 269)_ = 12.1; Saline, n = 22, N = 4; Psilocybin, n = 17; N = 6). **(B)** Reduced frequency of spontaneous EPSC of excitatory layer 5 pyramidal neurons of Psilocybin mice (*p*, Wilcoxon test; Saline, n = 22, N = 4; Psilocybin, n = 20; N = 5). **(C)** SST+ neurons didn’t show change of intrinsic excitability (Saline, n = 28, N = 3; Psilocybin, n = 20; N = 3). **(D)** F-I plot showed enhanced excitability of inhibitory PV+ neurons (Two-way ANOVA, F_(3, 332)_ = 5.6; Saline, n = 18, N = 5; Psilocybin, n = 30; N = 7). **(E** and **F**) PV+ neurons showed intrinsic properties changes. The threshold (**E**) and half-width **(F)** of action potential decreased (*p*, Wilcoxon test; Saline, n = 25, N = 6; Psilocybin, n = 30; N = 7). **(G)** Increased amplitude of spontaneous IPSC of PV+ neurons of Psilocybin mice (*p*, Wilcoxon test; Saline, n = 14, N = 6; Psilocybin, n = 12; N = 6).

For the GABAergic SST+ neurons, we observed no difference in intrinsic properties or synaptic inputs of SST+ neurons between Saline and Psilocybin mice (**Figure 3C**, **Figure S16C**, and **Figure S17C**).

Last, we recorded GABAergic PV+ neurons; the firing of these neurons increased in Psilocybin mice, with the threshold and half-width of action potential decreased (**Figure 3D** – **F** and **Figure S16B**). Interestingly, while the excitatory transmission did not change, the amplitude of sIPSC increased (**Figure 3G**, **Figure S16B**, and **Figure S17B**).

Taken together, given that layer 5 pyramidal neurons are the major output neurons of the cortex, these data suggest that psilocybin induced reduced activity in the OFC. Considering layer 5 pyramidal neurons are the major output neurons in the neocortex, these data suggest psilocybin reduced the output of the OFC.

### Cell-type-specific deletion of *Htr2a* in the OFC compromised the anti-depressant effect of psilocybin

Neurons in the brain interconnect and form circuits to exert functions. Neurons in circuits not only show plasticity with external stimuli, they also exhibit “homeostatic plasticity” that acts to stabilize neuronal and circuit activity to counterbalance activity changes of other neurons ^49^. To understand the mechanism of cell-type specific changes of neuronal activities in the OFC, we performed cell-type specific deletion of the *Htr2a* in neurons of the OFC and examined the protein expression of these neurons. By AAV-mediated Cre-dependent shRNA, we found that sh- *Htr2a* reduced the expression of 5-HT_2A_R in the infected neurons (**Figure 4A**). We then evaluated sh-*Htr2a* on GluR1 (*Gria1*) expression in layer 5 excitatory neurons, and found AAV-mediated deletion of *Htr2a* in the deep layer excitatory neurons abolished the reduction of GluR1 after psilocybin injection, suggesting such a reduction of GluR1 in excitatory neurons in the OFC is a direct effect of the activation of 5-HT_2A_R by psilocybin, but not via homeostatic regulation of the local circuit (**Figure 4B**).

**Figure 4.**
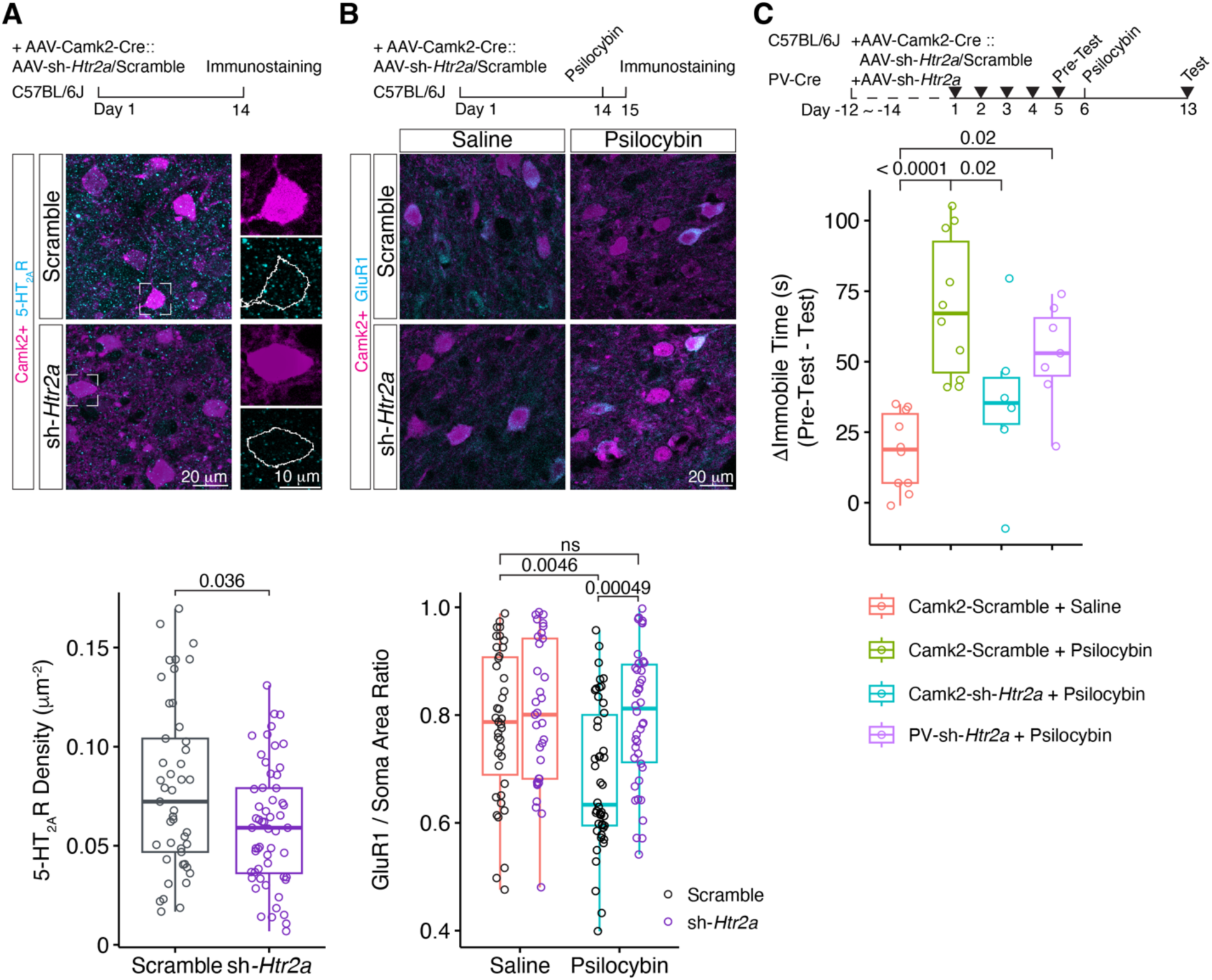
Knockdown of *Htr2a* blocked psilocybin induced GluR1 down-regulation and anti-depressant effect in rFST. **(A)** shRNA reduced 5-HT_2A_ receptor subunit expression in layer 5 pyramidal neurons (*p*, Wilcoxon test; Scramble, n = 45; sh-*Htr2a*, n = 56; N = 3/group). **(B)** sh-*Htr2a* blocked psilocybin induced GluR1 down-regulation in layer 5 pyramidal neurons. Psilocybin or saline was *i.p.* injected 24 hrs before tissue collection (Two-way ANOVA, F_(3,156)_ = 5.04, p = 0.026; *post hoc* Tukey test: Saline + Scramble vs Psilocybin + Scramble, *p* = 0.0046; Psilocybin + Scramble vs Psilocybin + sh-*Htr2a*, *p* = 0.00049; Saline + sh-*Htr2a* vs Psilocybin + Scramble, *p* = 0.00084; Saline + Scramble, n = 39; Saline + sh-*Htr2a*, n = 33; Psilocybin + Scramble, n = 46; Psilocybin + sh-*Htr2a*, n = 42; N = 3/group) **(C)** Cell-type specific knockdown of *Htr2a* on the anti-depressant effect of psilocybin in rFST. Top panel, experiment timeline, arrowheads indicate timepoints of FSTs. Psilocybin or saline was *<u>i.p.</u>* injected 24 hrs after the FST on day 5 (Pre-Test). Bottom panel, the comparisons of the difference in immobile times of the last two FSTs, namely FST on day 13 (Test) and FST on day 5 (Pre-Test) (One-way ANOVA, F_(3,29)_ = 10.09, *p* = 0.0001; *post hoc* Holm test: Camk2-Scramble + Saline vs Camk2-Scramble + Psilocybin, *p* = 6.3 x 10^-5^; Camk2-Scramble + Psilocybin vs Camk2-sh-*Htr2a* + Psilocybin, *p* = 0.02; Camk2-Scramble + Saline vs PV-sh-*Htr2a* + Psilocybin, *p* = 0.02; Camk2-Scramble + Saline, N = 10; Camk2-Scramble + Psilocybin, N = 10; Camk2-sh-*Htr2a* + Psilocybin, N= 6; PV-sh-*Htr2a* + Psilocybin, N = 7).

Recent studies have shown the therapeutic potential of psilocybin as an anti-depressant. A single dose of psilocybin promoted a quick and long-lasting anti-depressant effect in both animal and clinical studies ^8, 21, 50–52^. To illustrate the contribution of the neurons in the OFC to the anti-depressant effect of psilocybin, we induced cell-type specific deletion of *Htr2a*, then evaluated the effect on animal behavior (**Figure 4C** and **S18**). We injected Cre-dependent sh-*Htr2a* into the OFC of mice and then evaluated the anti-depressant effect of psilocybin with a chronic stress model, specifically, repeated forced swimming, which induces a depressed state ^52–54^ (**Figure 4C**). Compared with mice injected with saline, a single injection of psilocybin reduced the immobile time of mice in the repeated forced swimming test (**Figure 4C**). Furthermore, knockdown of *Htr2a* in excitatory neurons in the OFC abated the anti-depressant effect of psilocybin, while knockdown of *Htr2a* in PV+ neurons only partially reduced the anti-depressant effect of psilocybin (**Figure 4C**).

## Discussion

In the present study, we combined single nucleus RNA-seq with functional assays to study the long-term effect of psilocybin on cells of the orbitofrontal cortex (OFC), especially on neurons. We showed psilocybin-induced gene expression changes in neurons of the OFC, which contribute to the functional changes in these neurons and the OFC. Furthermore, such function changes depending on the expression of 5-HT_2A_ receptors in excitatory neurons of the deep layers as knockdown of *Htr2a* reversed psilocybin-induced changes in GluR1 (*Gria1*) and its anti-depressant effect.

Our recordings from layer 5 pyramidal excitatory neurons, inhibitory PV+, and SST+ neurons showed that psilocybin induced different neuronal activity changes in these three types of neurons. In the agranular cortex, including prefrontal cortical regions and the motor cortex, studies have shown that in a microcircuit composed of these three types of neurons, while the pyramidal neurons serve as the main output, the tone of inhibition to these neurons is determined by PV+ neuron activity with SST+ neurons mainly modulate PV+ neuron activity ^55, 56^. In the present study, we observed decreased synaptic transmission and output from layer 5 pyramidal neurons and increased output from PV+ neurons, while those of the SST+ neurons did not change after psilocybin injection. Our findings indicate that psilocybin led to decreased network activity in the OFC. Such a reduction is also consistent with previous studies, which showed that activation of 5-HT_2A_R in the OFC decreased population neuronal activity ^57^.

Our snRNA-seq results showed that the Ex1 cluster of the OFC neurons showed the most significant gene expression changes in Psilocybin mice. Besides reduced expression of glutamate receptors, the most affected genes are involved in synaptic transmission, cell-adhesion at synapse, and synapse formation and maintenance. Analysis of those cell-cell interaction signaling pathways showed that besides the Epha5-Efna5 signaling pathway, other affected pathways between neuronal clusters all majorly showed reductions in Psilocybin mice. Meanwhile, it is interesting that we did not observe changes in the expression of genes involved in excitatory synaptic transmission or recorded changes in sEPSCs in both PV+ and SST+ neurons. Furthermore, deletion of *Htr2a* in PV+ neurons did not abolish the anti-depressant effect of psilocybin. These data suggest that these two types of interneurons might not be the main target of psilocybin, which in part supports the view that deep-layer pyramidal neurons are important for the effects of psilocybin. It is unclear whether the relatively low expression of *Htr2a* in these neurons contributes to their relative insensitivity to psilocybin.

Psilocybin is metabolized into psilocin in the body, which is an agonist of 5-HT_2A_ receptor (5-HT_2A_R). 5-HT_2A_Rs are G-protein coupled receptors, which activates intracellular G_αq_-like G proteins and β-arrestin signaling pathway. While the downstream targets of the activation of 5-HT_2A_R is not fully illustrated, studies have shown that psilocybin and other psychedelics induced long-term plasticity change and gene expression change in neurons ^58^, suggesting increased excitatory transmission in the brain. On the other hand, studies also showed that lesion or chronic inactivation of the OFC decreased depression-like behavior in rats ^59, 60^.

The orbitofrontal cortex is well known for its role in decision making and is essential for encoding information about rewards ^61^. Recent studies also showed that the OFC is vulnerable in psychological disorders. For example, clinical studies have shown that in both adults and adolescents with MDD the OFC were one of the most affected brain regions, showing thinner cortical gray matter or reduced total surface area ^62^. Furthermore, the effectiveness of the antidepressant treatment is associated with the decrease of the activation of the OFC ^29^, and anti-depressants on healthy subjects reduced connectivity of the OFC with subcortical regions, such as the amygdala and the striatum ^63^. The OFC connect with many cortical and subcortical regions, among its target both the ventral medial prefrontal cortex and the posterior cingulate cortex are also major components of the default-mode network (DMN) ^26^. The DMN exists not only in human, also in primates and rodents ^64, 65^. Studies implicated depressive symptomology associated with hyperconnectivity of the DMN ^23^, increased functional connectivity of the posterior cingulate cortex with the OFC in MDD patients, and anti-depressants also reduced this functional connectivity ^28^. Furthermore, in the brain, psilocybin and other psychedelics reduced the activity and functional connectivity of the DMN ^25, 66^, while single dose of psilocybin also decreased DMN recruitment in patients with treatment-resistant depression ^24^. Taken together, these studies have suggested that the anti-depressant effect of psychedelics and traditional anti-depressants is associated with reduced activity and/or output of the OFC.

In conclusion, by combining snRNA-seq with electrophysiology and behavior tests, we showed that psilocybin induced cell-type-specific long-term changes in the neurons of the OFC. These changes converged to decrease circuit activity of the OFC, and contributed to the anti-depressant effect of psilocybin.

## Acknowledgements

Research was supported in part by Ministry of Science and Technology of China (STI2030-Major Projects 2021ZD0202900 and 2019YFA0706201, W.Z.), National Natural Science Foundation of China (32170960, W.Z.).

## Author contributions

W.Z. conceived and designed the experiments. Z.H., X.W., Y.W., and performed the experiments. Z.H., X.W., Y.W., J.T., J.D, B.L., and W.Z. analyzed and interpreted the data. W.Z. and L.B. wrote the manuscript with inputs from other authors.

---

**Figure S1.**
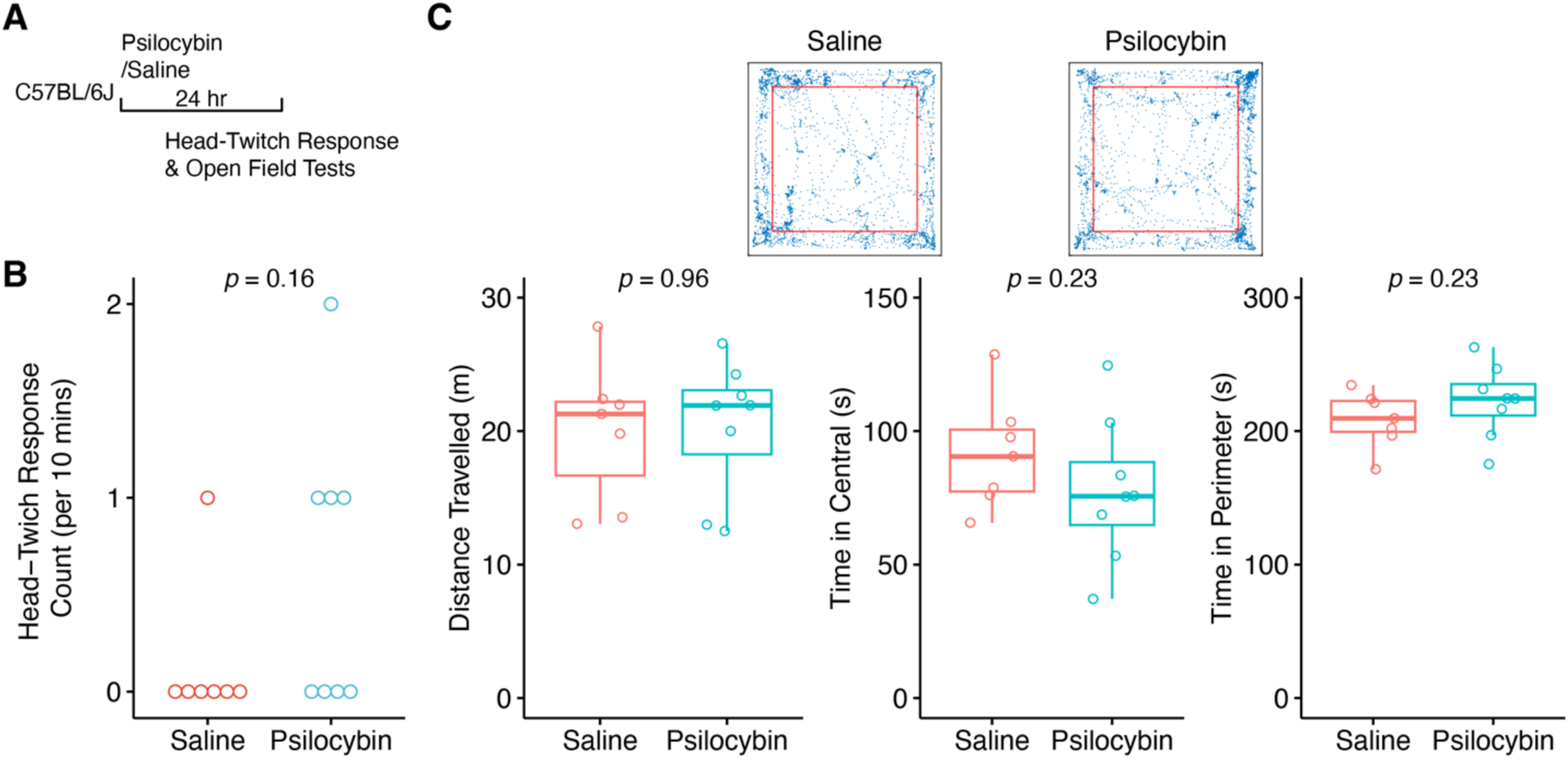
Psilocybin mice did not show behavior difference 24 hrs after injection compared with Saline mice, related to Figure 1. **(A)** Experiment timeline. **(B)** Head-twitch response counts in 10 mins of mice 24 hrs after *i.p.* injection. **(C)** Open field test of mice 24 hrs after injection. Top panels, representative traces of mice in open field arena for 5 mins with red squares denote the central area. Bottom panels, statistical comparisons of the distance mice travelled in the arena (left panel), time in the central area (middle panel), and time in the perimeter area (right panel) for 5 mins. *P*, Wilcoxon test; Saline, N = 7; Psilocybin, N = 8. Circles indicate individual measurements.

**Fig S2.**
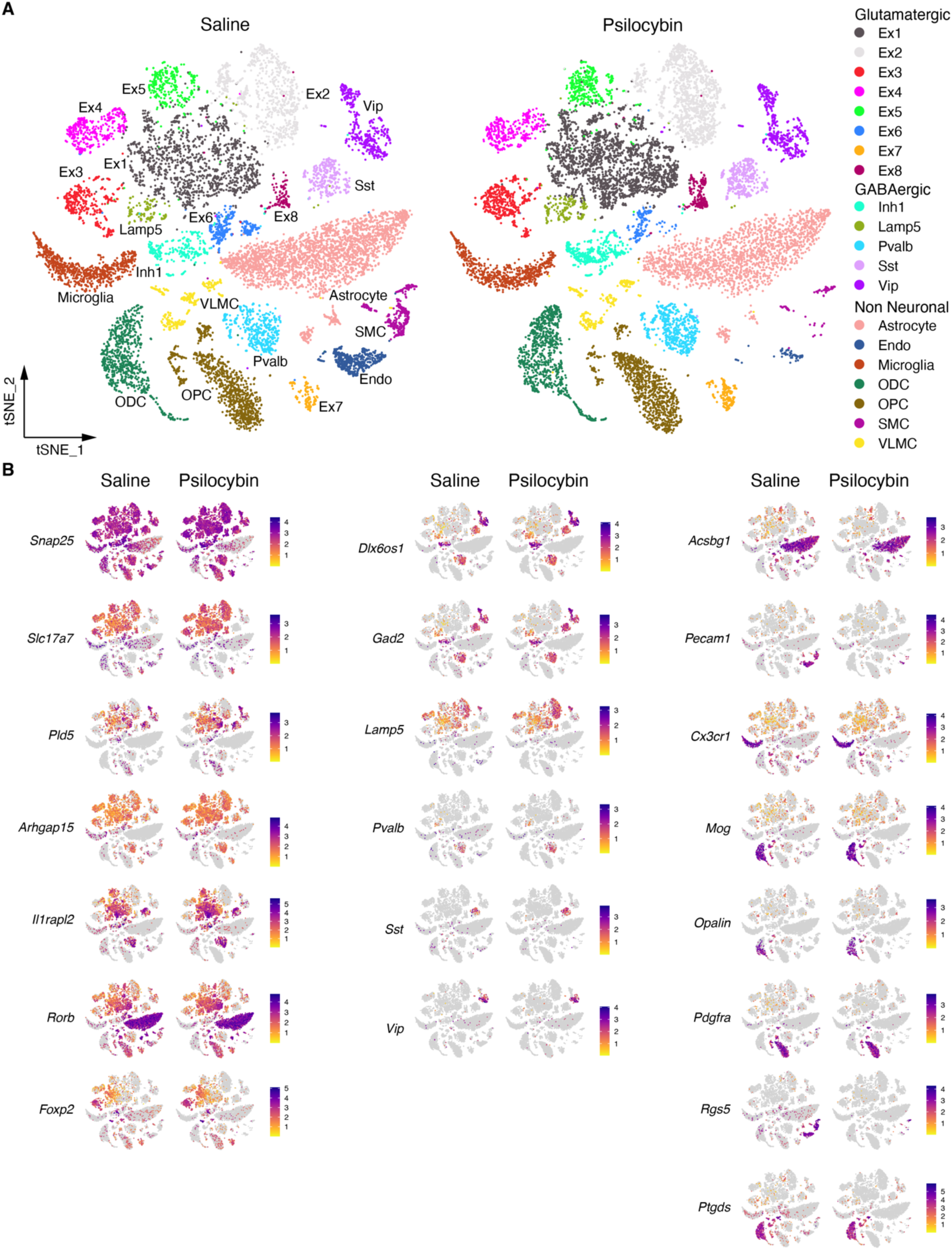
The expression of cell-type markers in the OFC, related to Figure 1. **(A)** *t*-SNE plots of cells from Saline and Psilocybin mice. Clusters colored based on cell-types. **(B)** *t*-SNE plots of the expression of the selected genes used for cell type identification. Saline, n = 12246; Psilocybin, n = 12991; N = 4/group.

**Figure S3.**
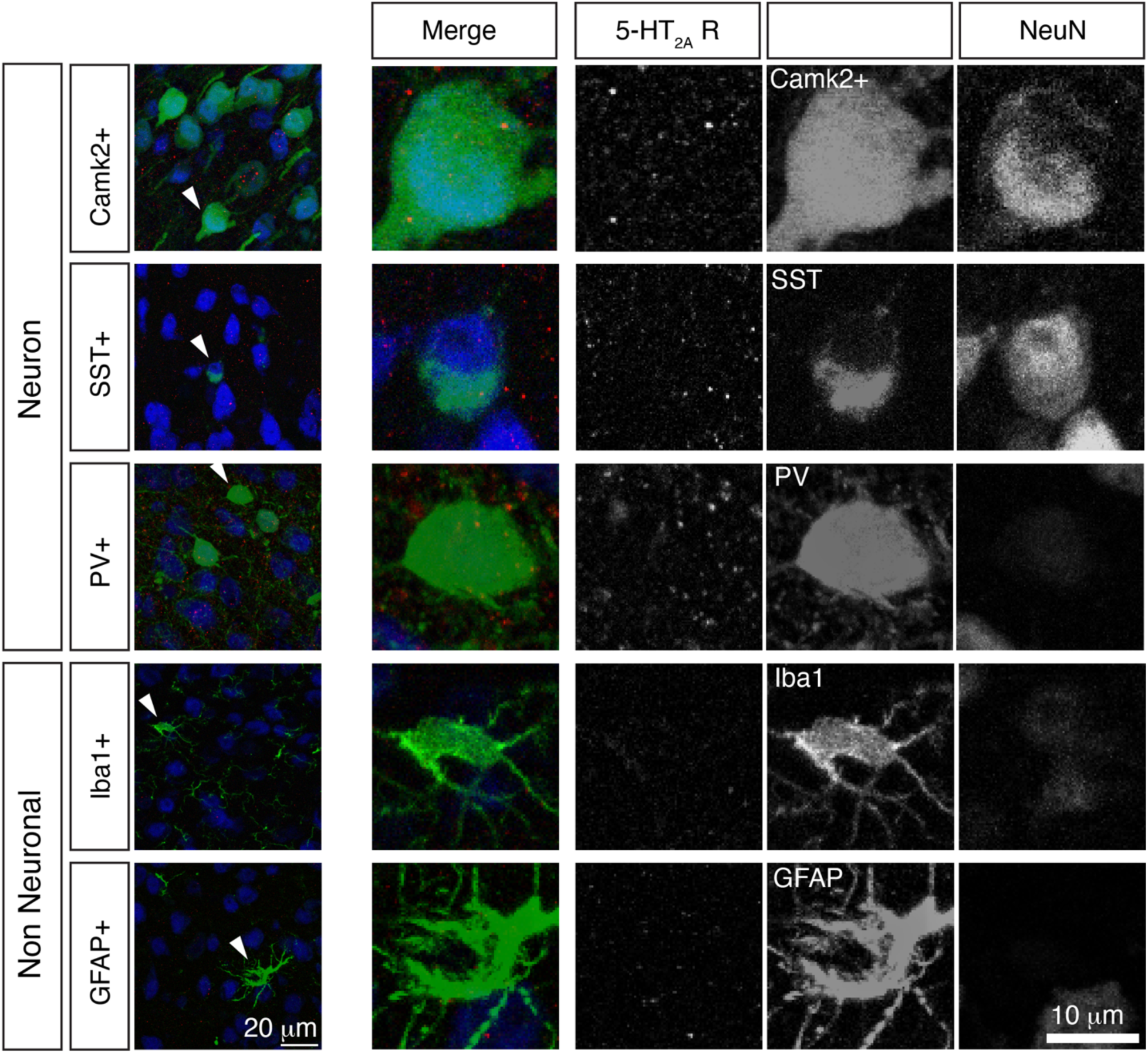
The expression of 5-HT_2A_ receptor in cells of the OFC, related to Figure 1. 5-HT_2A_ receptor subunit expresses neurons (CamkII+, SST+, and PV+) but not microglia (Iba1+) or astrocyte (GFAP+) cells. Arrowheads in the left panel indicate the cells showed on the right.

**Figure S4.**
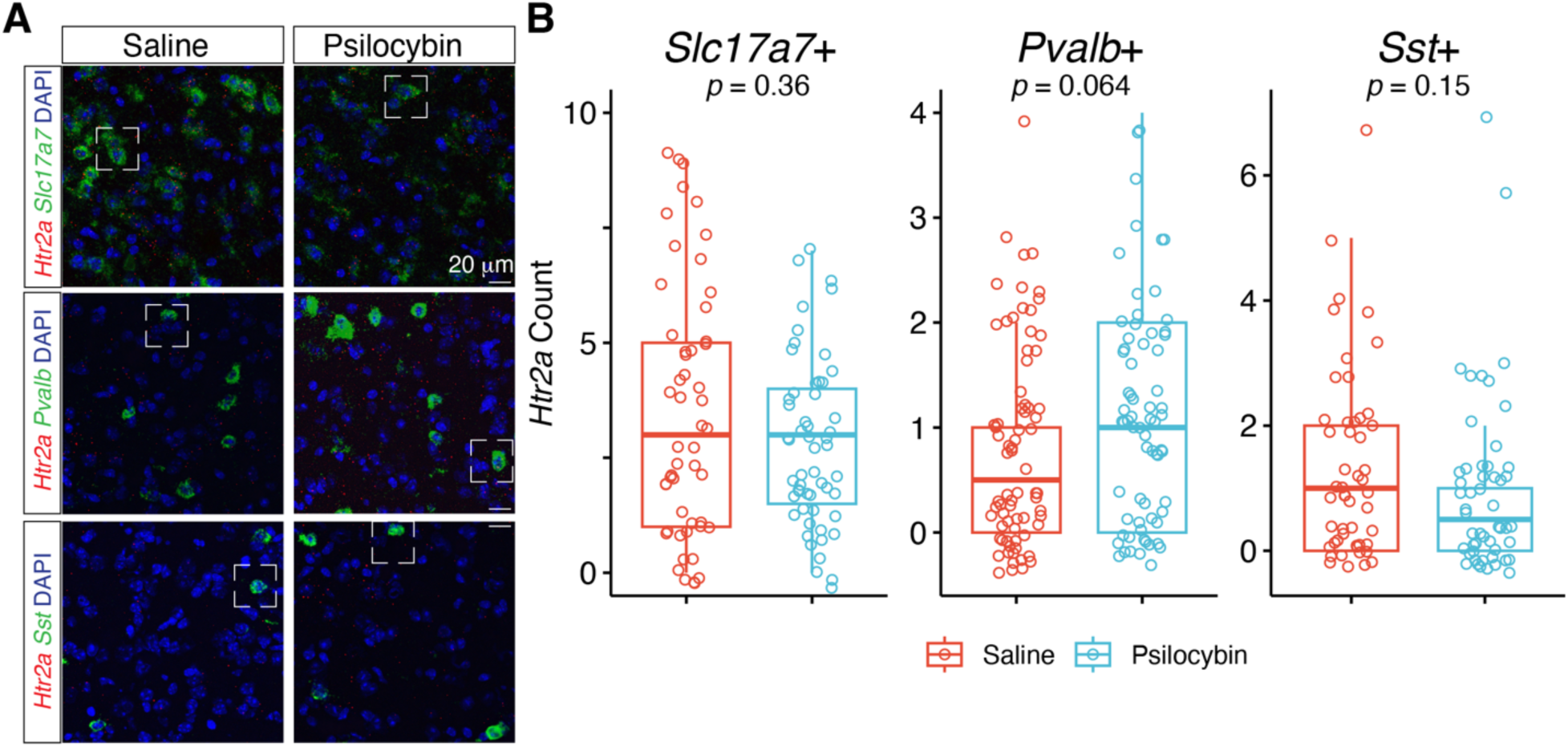
Statistical analysis of RNAscope *in situ* hybridization of *Htr2a* mRNA in cells of the OFC, related to Figure 1. **(A)** Representative images of *Htr2a* mRNA expression in excitatory (*Slc17a7*+), inhibitory PV+ (*Pvalb*+), and inhibitory SST+ (*Sst*+) neurons of layer 5 in the OFC. Dashed squares indicate the cells shown in Figure **1E**. **(B)** Statistical comparison of *Htr2a* puncta counts in *Slc17a7*+, *Pvalb*+, and *Sst*+ cells (*p*, Wilcoxon test; *Slc17a7*+: Saline, n =52; Psilocybin, n = 55; *Pvalb*+: Saline, n = 74; Psilocybin, n = 68; *Sst*+: Saline, n = 50; Psilocybin, n = 52; N = 3/group).

**Figure S5.**
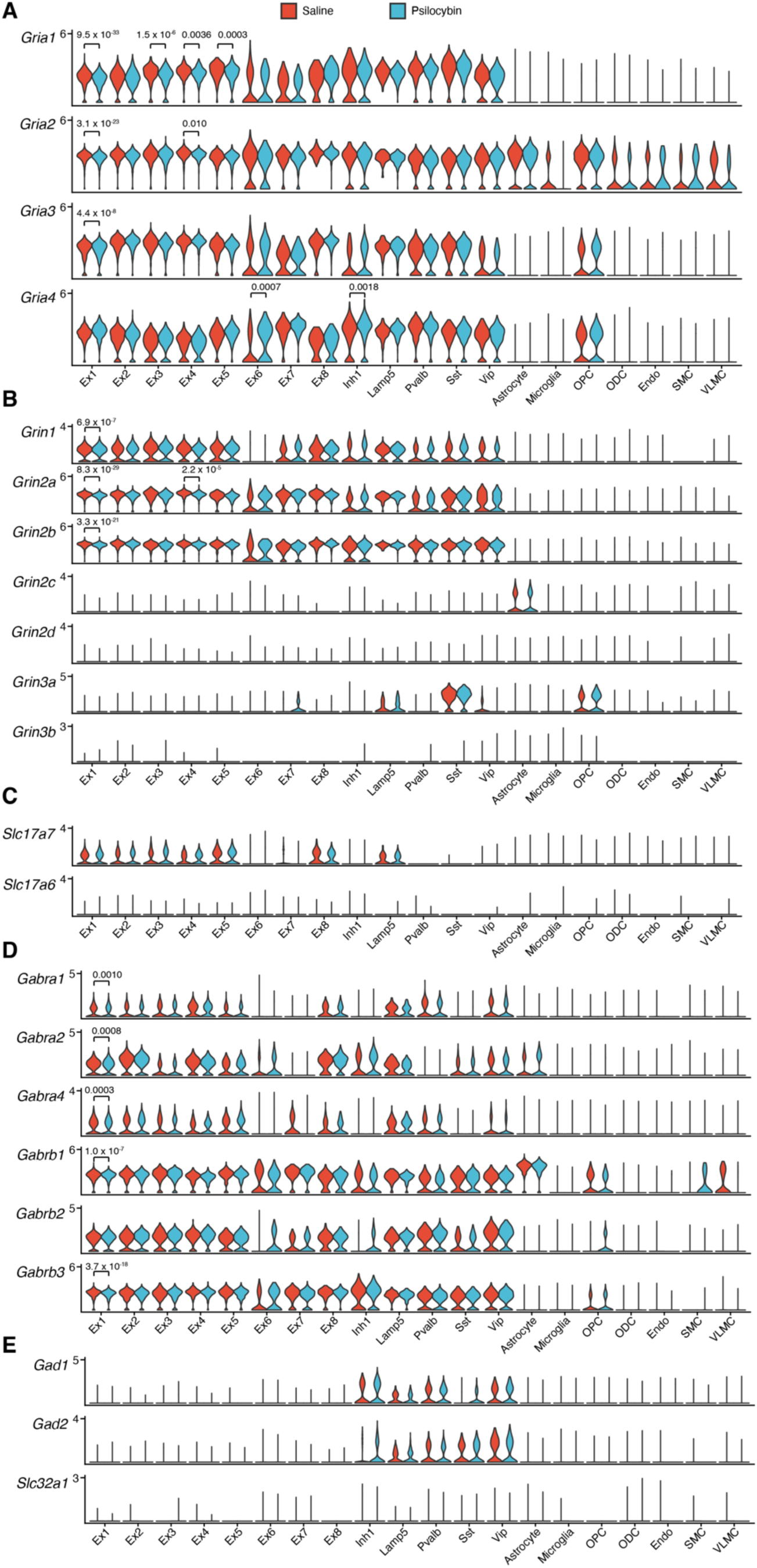
Comparisons of gene expressions of molecules involved in the excitatory and inhibitory synaptic transmission in clusters of the OFC between Saline and Psilocybin mice, related to Figure 1. **(A– C)** The comparisons of mRNA expression of molecules in the excitatory synaptic transmission, including AMPA receptor subunits **(A)**, NMDA receptor subunits **(B)**, and vesicular glutamate transporters **(C)**. **(D** – **E)** The comparisons of mRNA expression of molecules in the inhibitory synaptic transmission, including major GABA_A_ receptor subunits **(D)** and GABA synthesis and transport **(E)**. *P*, Wald test with Benjamini-Hochberg procedure, calculated with DESeq2.

**Figure S6.**
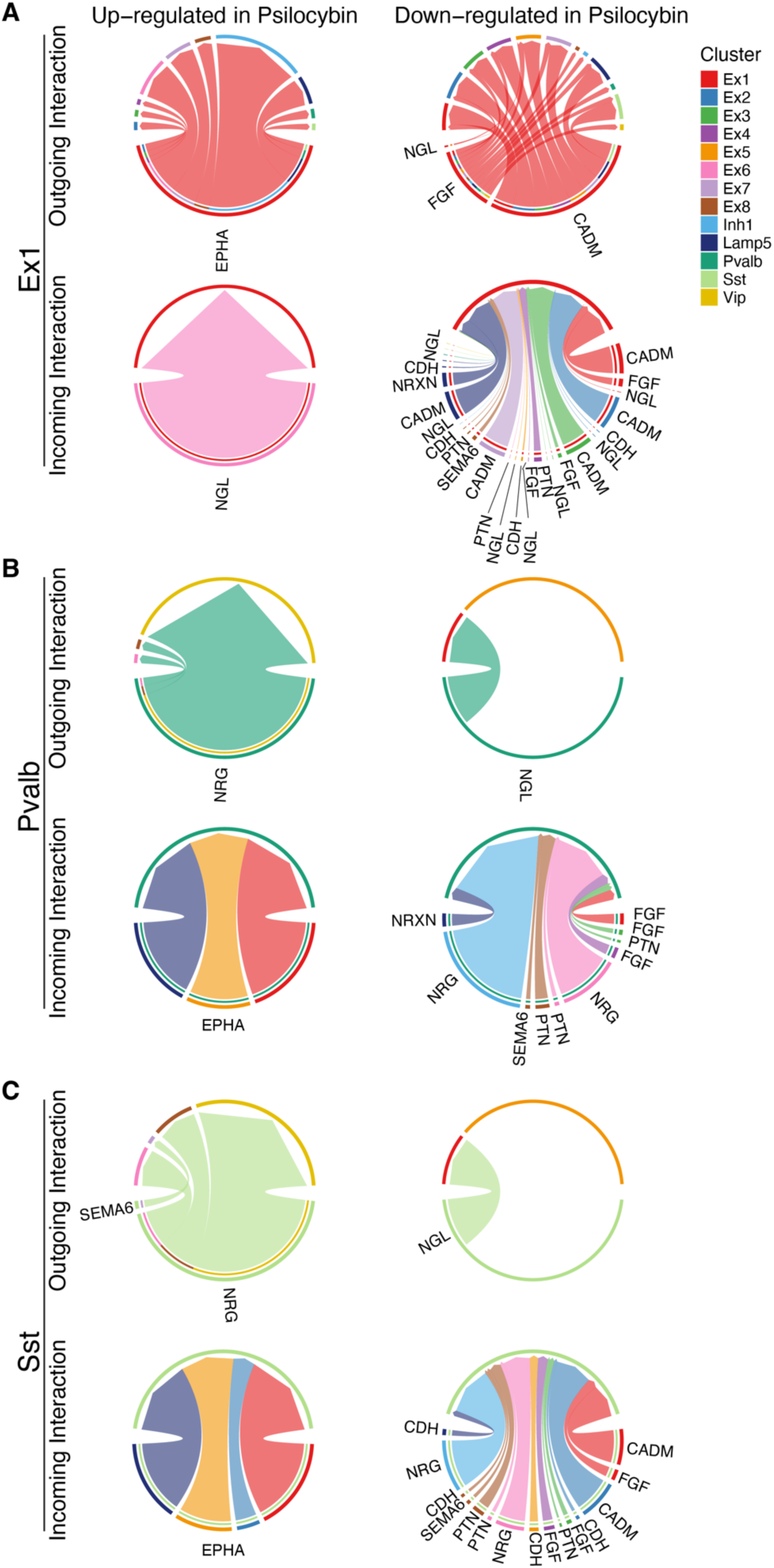
The cell-cell interaction signaling pathways of the Ex1, Pvalb, Sst clusters affected in Psilocybin mice, related to Figure 2. Chord diagrams show the changed cell-cell interaction of Ex1 (A), Pvalb (B), Sst (C) clusters with other neuronal cell clusters of Psilocybin mice, compared with Saline mice.

**Figure S7.**
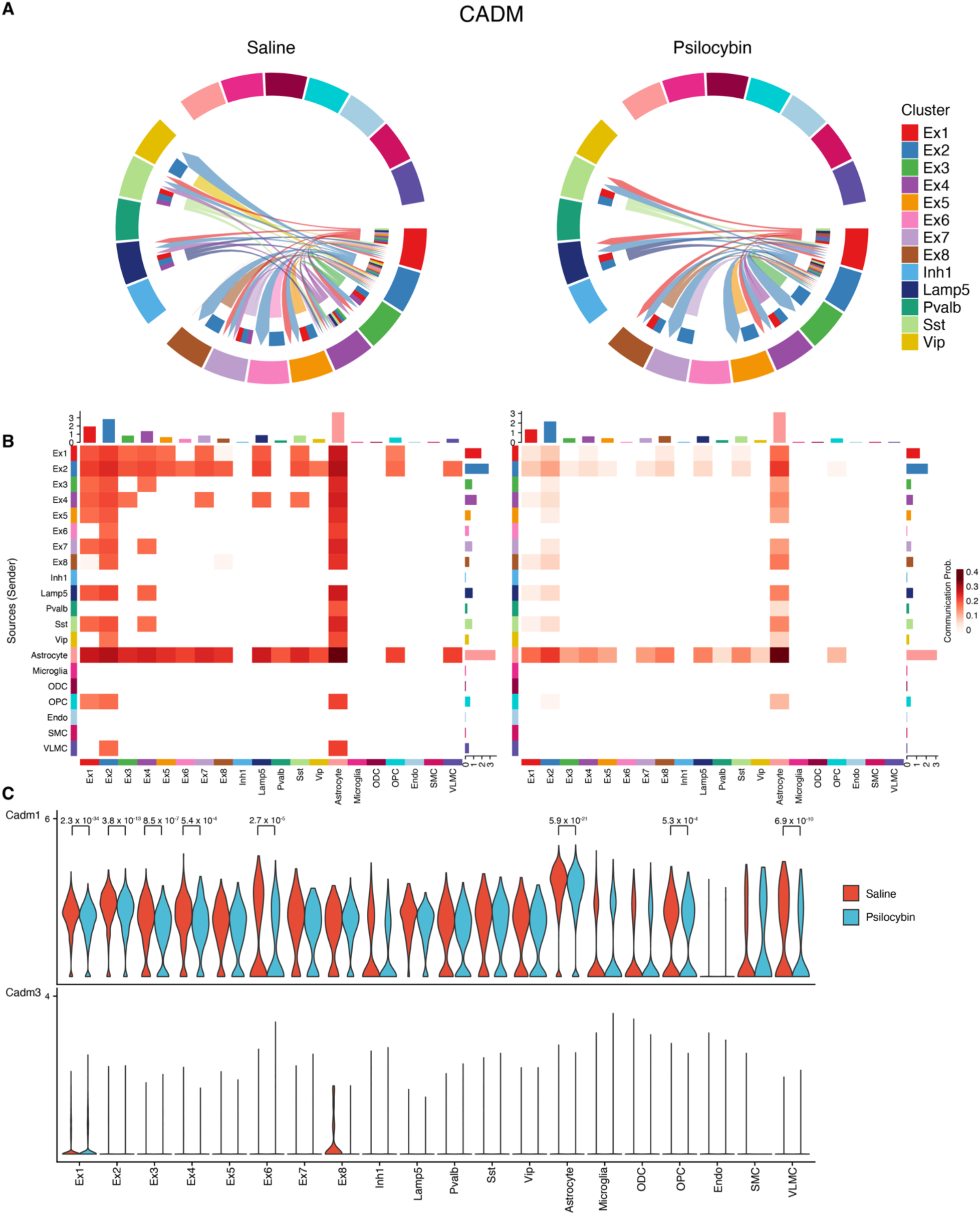
The comparison of the CADM signaling pathway of clusters between Saline and Psilocybin mice, related to Figure 2. **(A)** Chord diagrams of the CADM signaling pathway interaction strength of neuron clusters of saline and psilocybin mice. **(B)** The heatmaps of the CADM signaling pathway communication probability between cell clusters of saline and psilocybin mice. **(C)** The comparison of the CADM signaling pathway-related gene expression distribution in clusters of saline and psilocybin mice. *P*, Wald test with Benjamini-Hochberg procedure, calculated with DESeq2.

**Figure S8.**
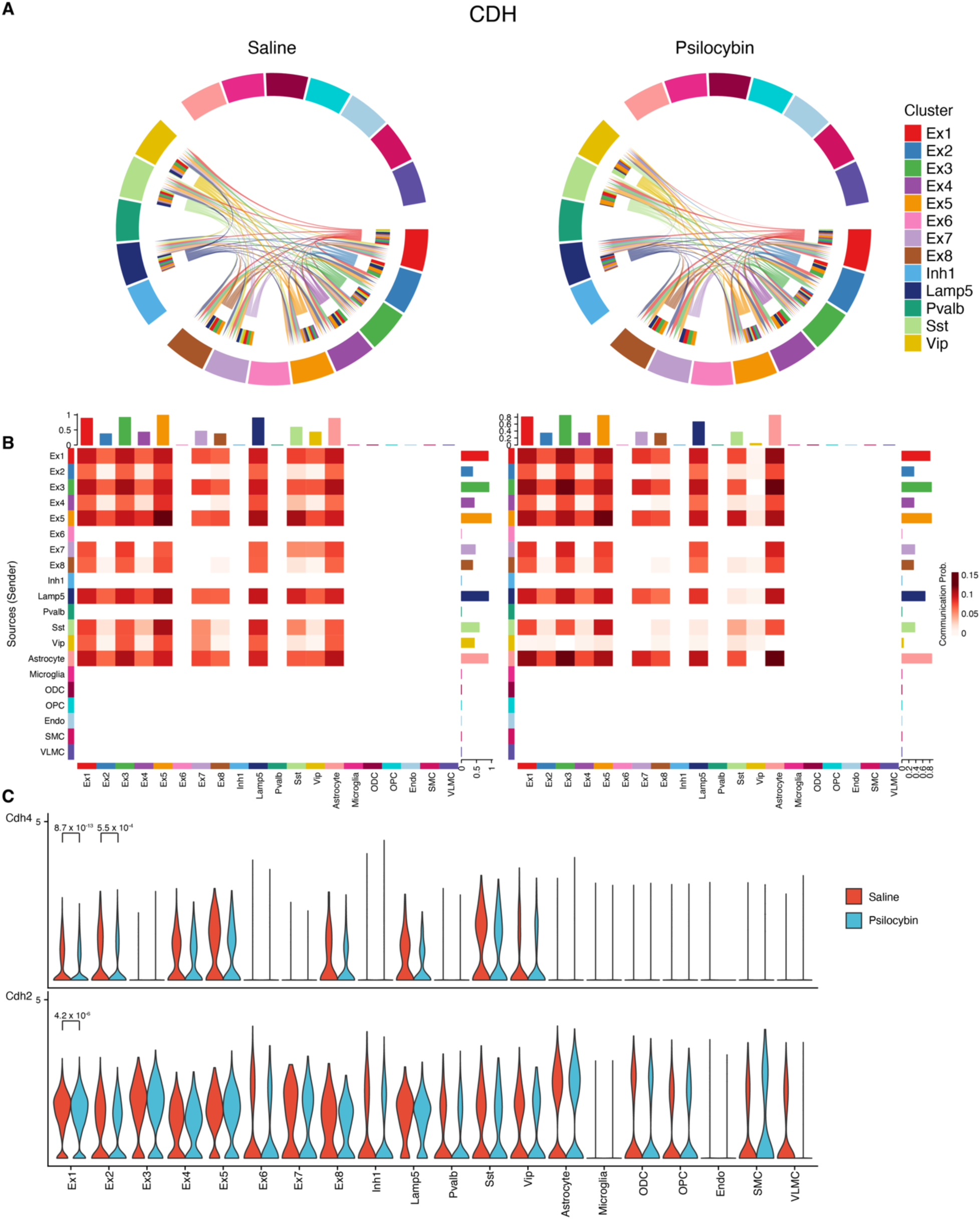
The comparison of the CDH signaling pathway of clusters between saline and psilocybin mice, related to Figure 2. **(A)** Chord diagrams of the CDH signaling pathway interaction strength of neuron clusters of saline and psilocybin mice. **(B)** The heatmaps of the CDH signaling pathway communication probability between cell clusters of saline and psilocybin mice. **(C)** The comparison of the CDH signaling pathway-related gene expression distribution in clusters of saline and psilocybin mice. *P*, Wald test with Benjamini-Hochberg procedure, calculated with DESeq2.

**Figure S9.**
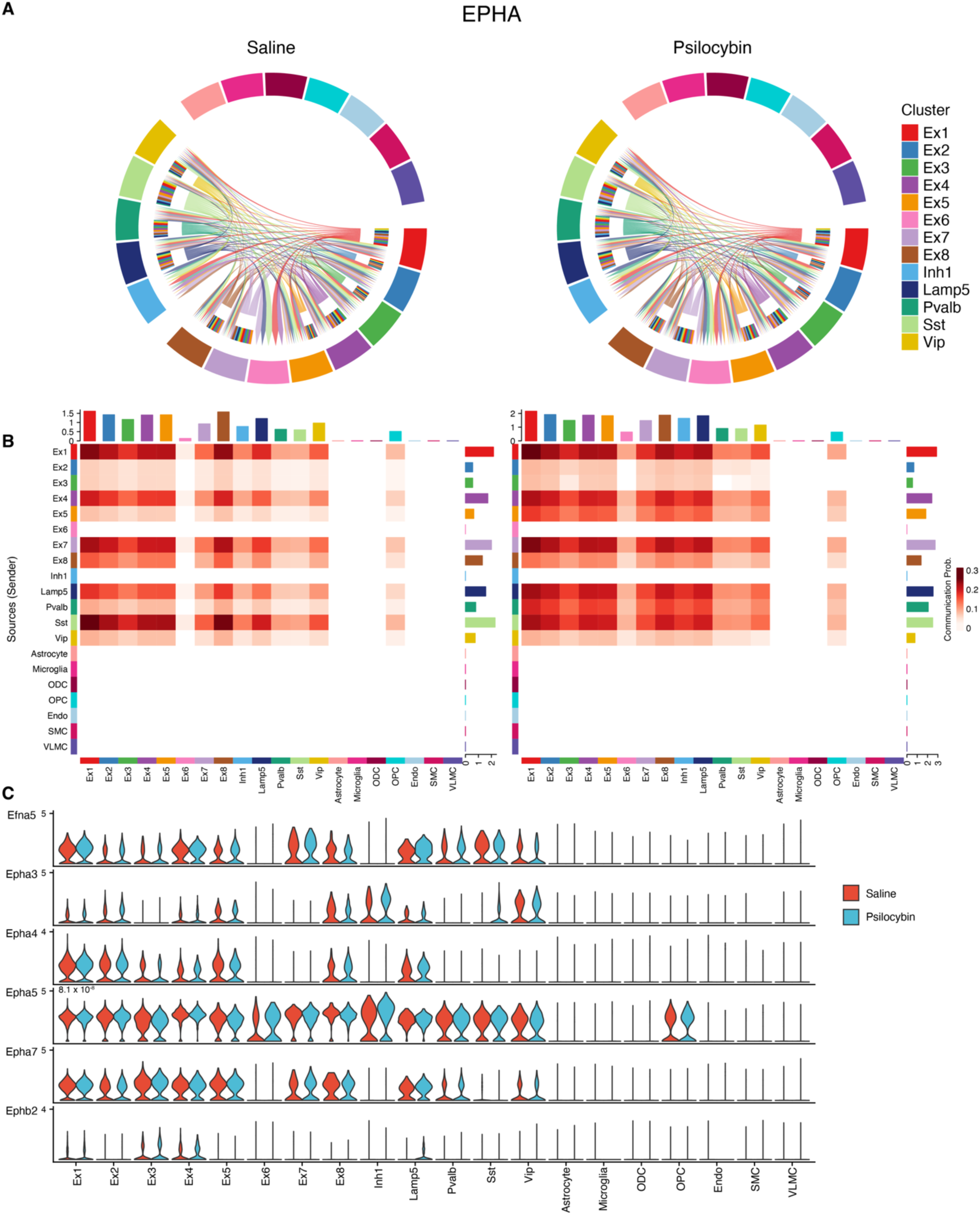
The comparison of the EPHA signaling pathway of clusters between saline and psilocybin mice, related to Figure 2. **(A)** Chord diagrams of the EPHA signaling pathway interaction strength of neuron clusters of saline and psilocybin mice. **(B)** The heatmaps of the EPHA signaling pathway communication probability between cell clusters of saline and psilocybin mice. **(C)** The comparison of the EPHA signaling pathway-related gene expression distribution in clusters of saline and psilocybin mice. *P*, Wald test with Benjamini-Hochberg procedure, calculated with DESeq2.

**Figure S10.**
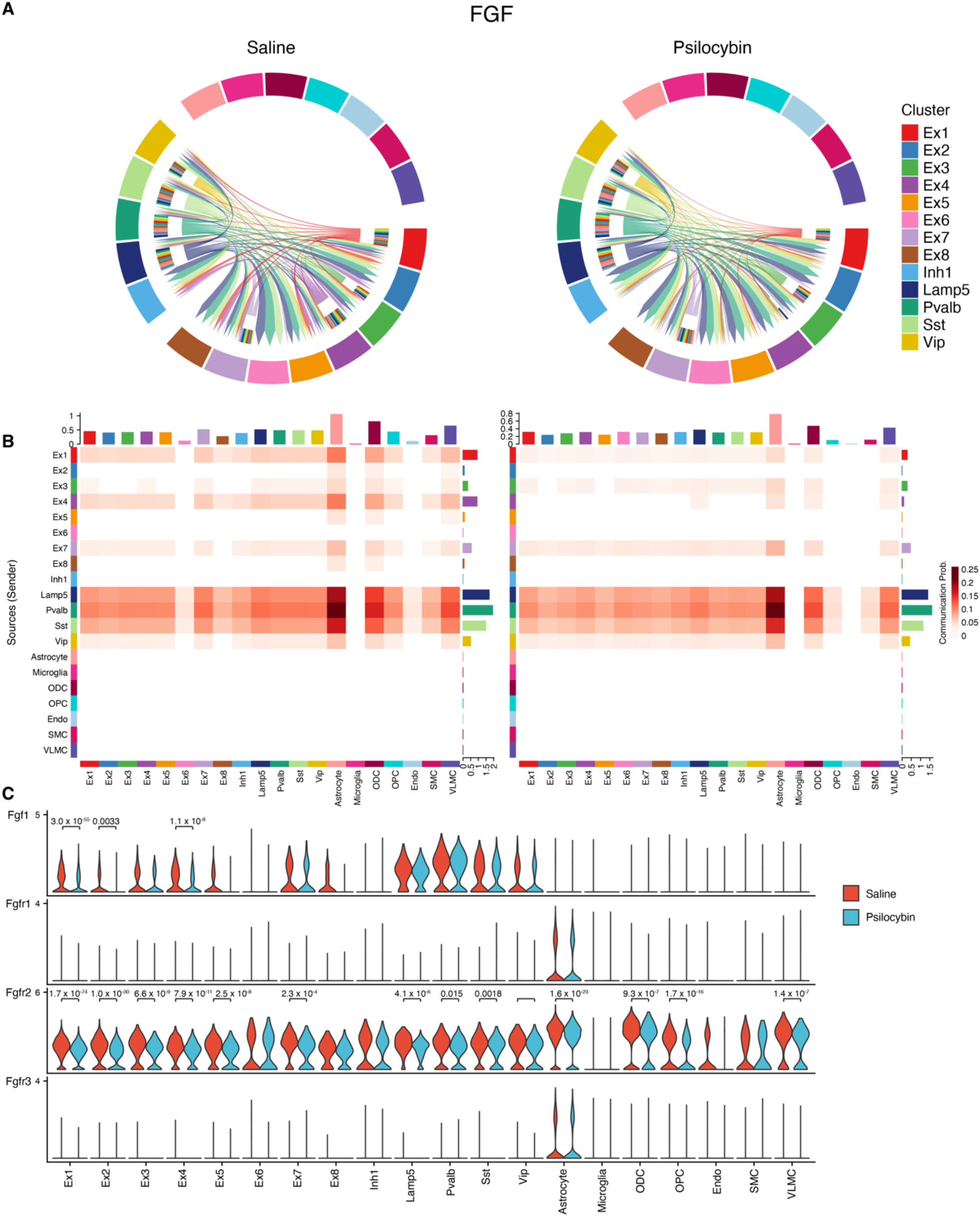
The comparison of the FGF signaling pathway of clusters between saline and psilocybin mice, related to Figure 2. **(A)** Chord diagrams of the FGF signaling pathway interaction strength of neuron clusters of saline and psilocybin mice. **(B)** The heatmaps of the FGF signaling pathway communication probability between cell clusters of saline and psilocybin mice. **(C)** The comparison of the FGF signaling pathway-related gene expression distribution in clusters of saline and psilocybin mice. *P*, Wald test with Benjamini-Hochberg procedure, calculated with DESeq2.

**Figure S11.**
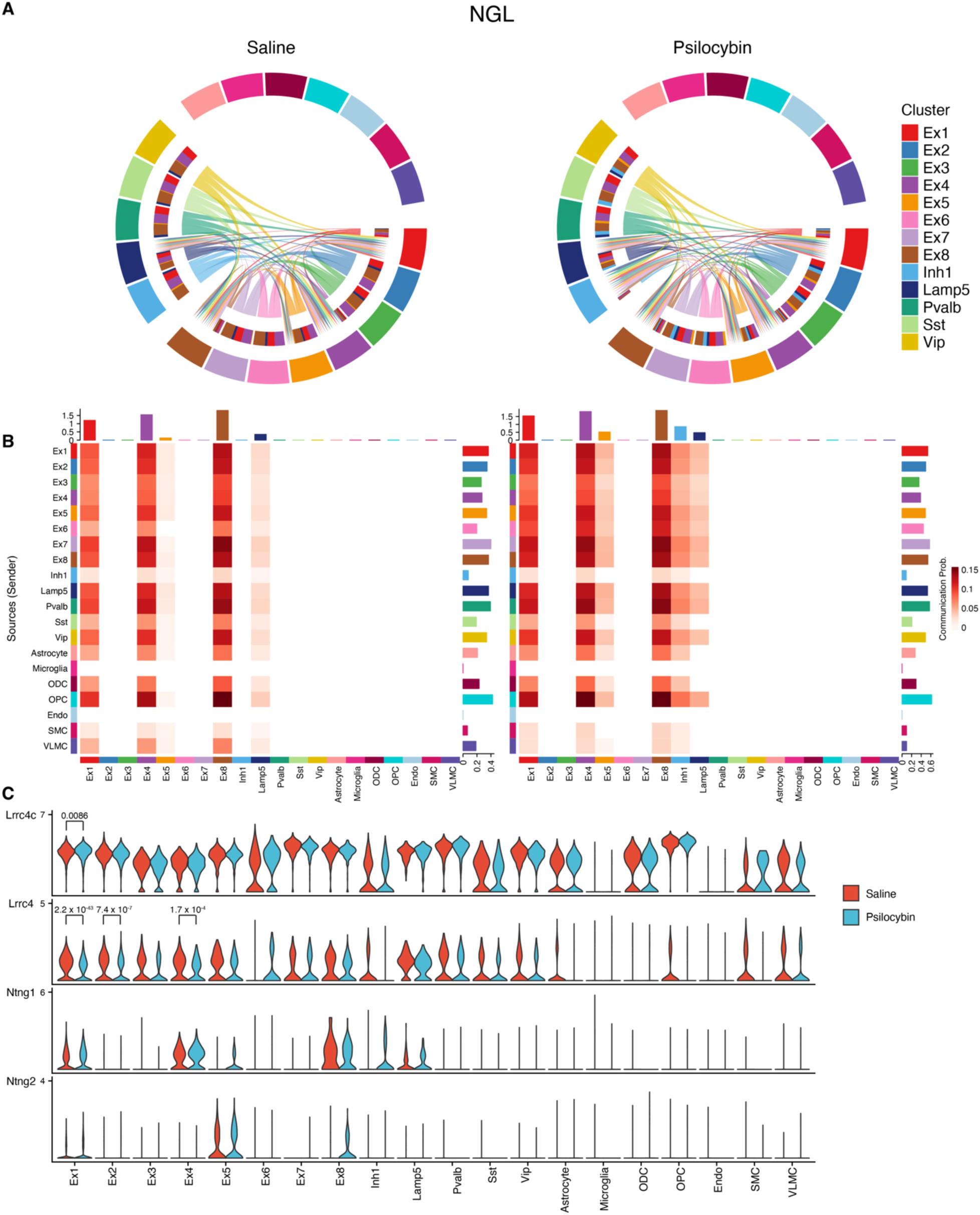
The comparison of the NGL signaling pathway of clusters between saline and psilocybin mice, related to Figure 2. **(A)** Chord diagrams of the NGL signaling pathway interaction strength of neuron clusters of saline and psilocybin mice. **(B)** The heatmaps of the NGL signaling pathway communication probability between cell clusters of saline and psilocybin mice. **(C)** The comparison of the NGL signaling pathway-related gene expression distribution in clusters of saline and psilocybin mice. *P*, Wald test with Benjamini-Hochberg procedure, calculated with DESeq2.

**Figure S12.**
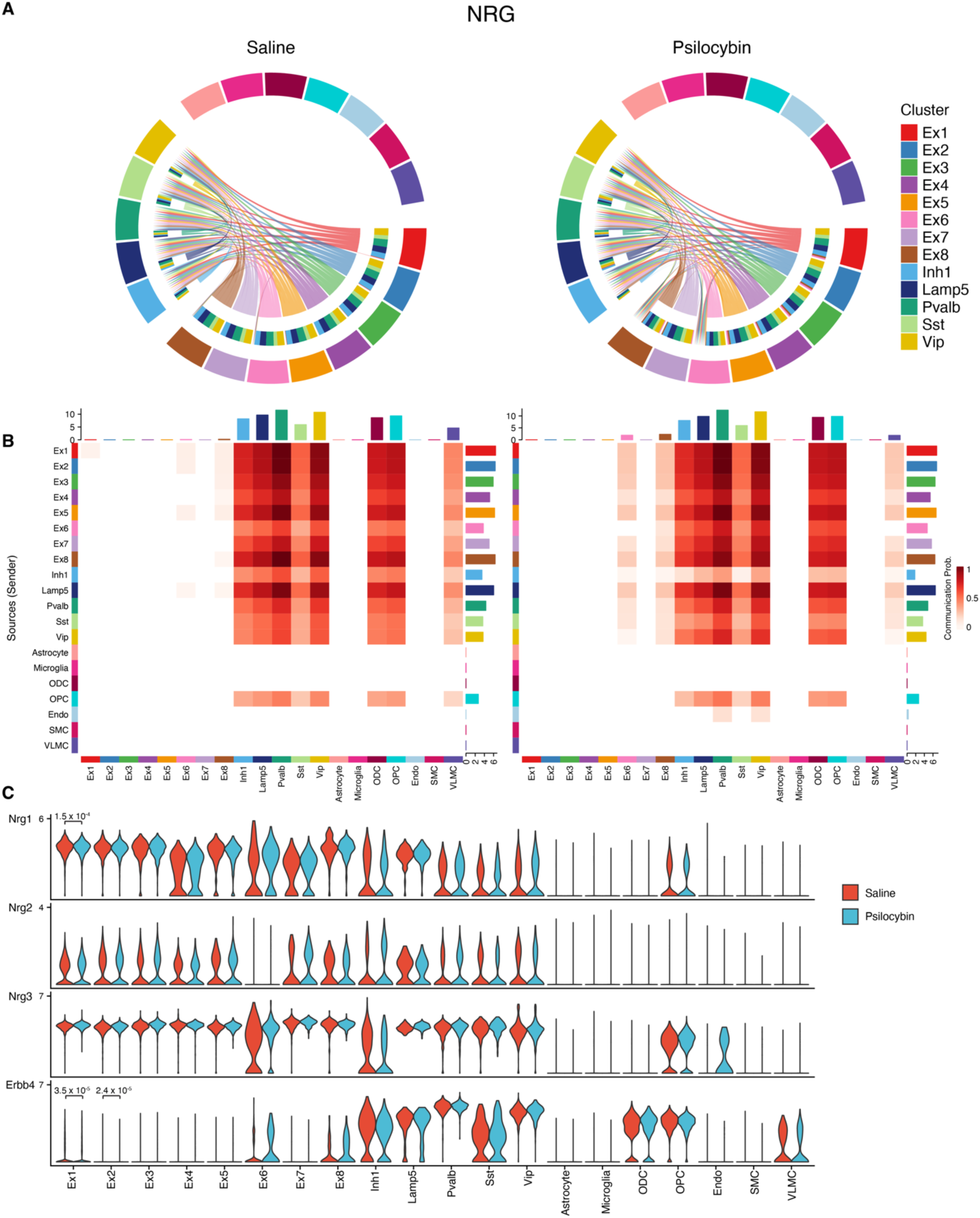
The comparison of the NRG signaling pathway of clusters between saline and psilocybin mice, related to Figure 2. **(A)** Chord diagrams of the NRG signaling pathway interaction strength of neuron clusters of saline and psilocybin mice. **(B)** The heatmaps of the NRG signaling pathway communication probability between cell clusters of saline and psilocybin mice. **(C)** The comparison of the NRG signaling pathway-related gene expression distribution in clusters of saline and psilocybin mice. *P*, Wald test with Benjamini-Hochberg procedure, calculated with DESeq2.

**Figure S13.**
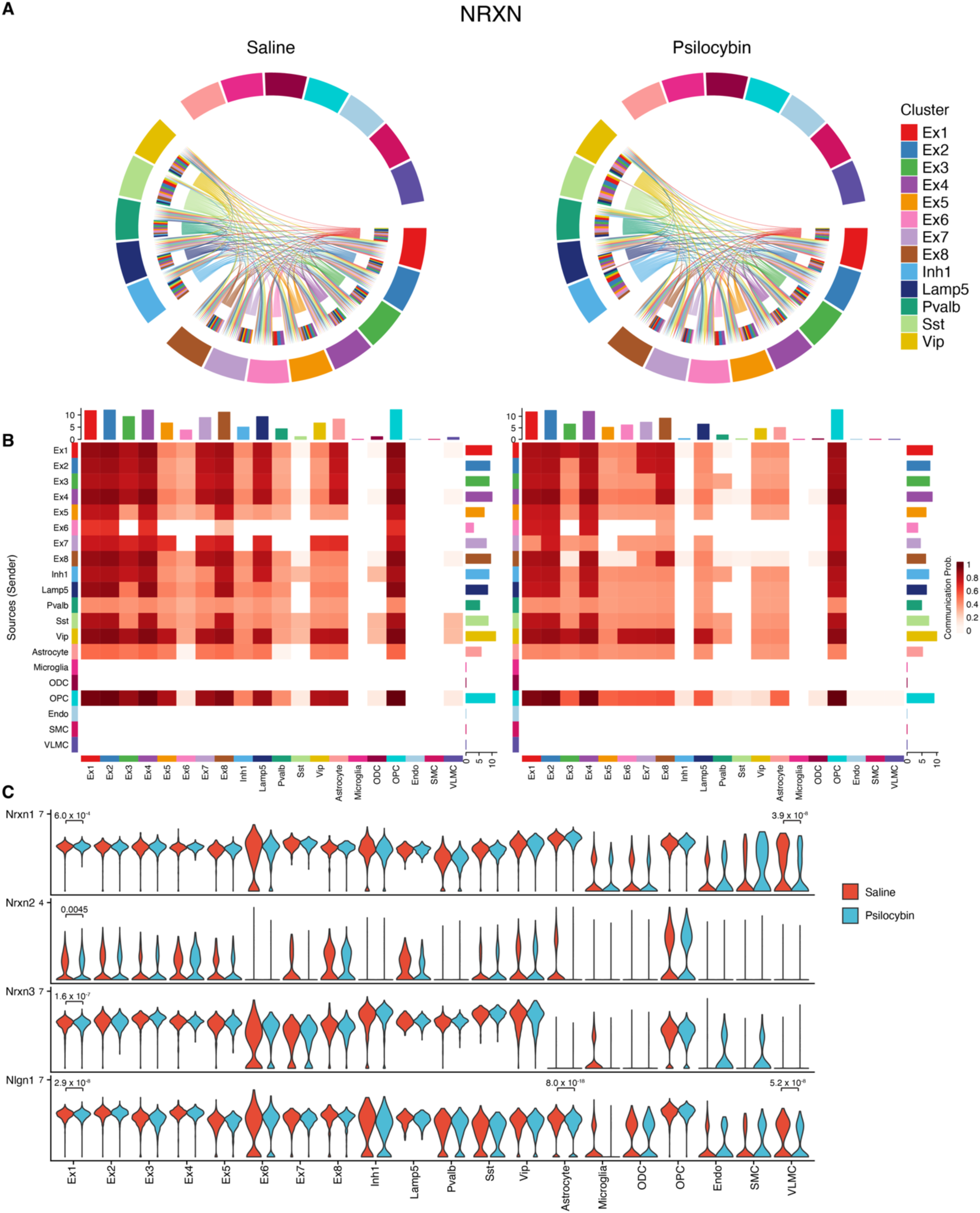
The comparison of the NRXN signaling pathway of clusters between saline and psilocybin mice, related to Figure 2. **(A)** Chord diagrams of the NRXN signaling pathway interaction strength of neuron clusters of saline and psilocybin mice. **(B)** The heatmaps of the NRXN signaling pathway communication probability between cell clusters of saline and psilocybin mice. **(C)** The comparison of the NRXN signaling pathway-related gene expression distribution in clusters of saline and psilocybin mice. *P*, Wald test with Benjamini-Hochberg procedure, calculated with DESeq2.

**Figure S14.**
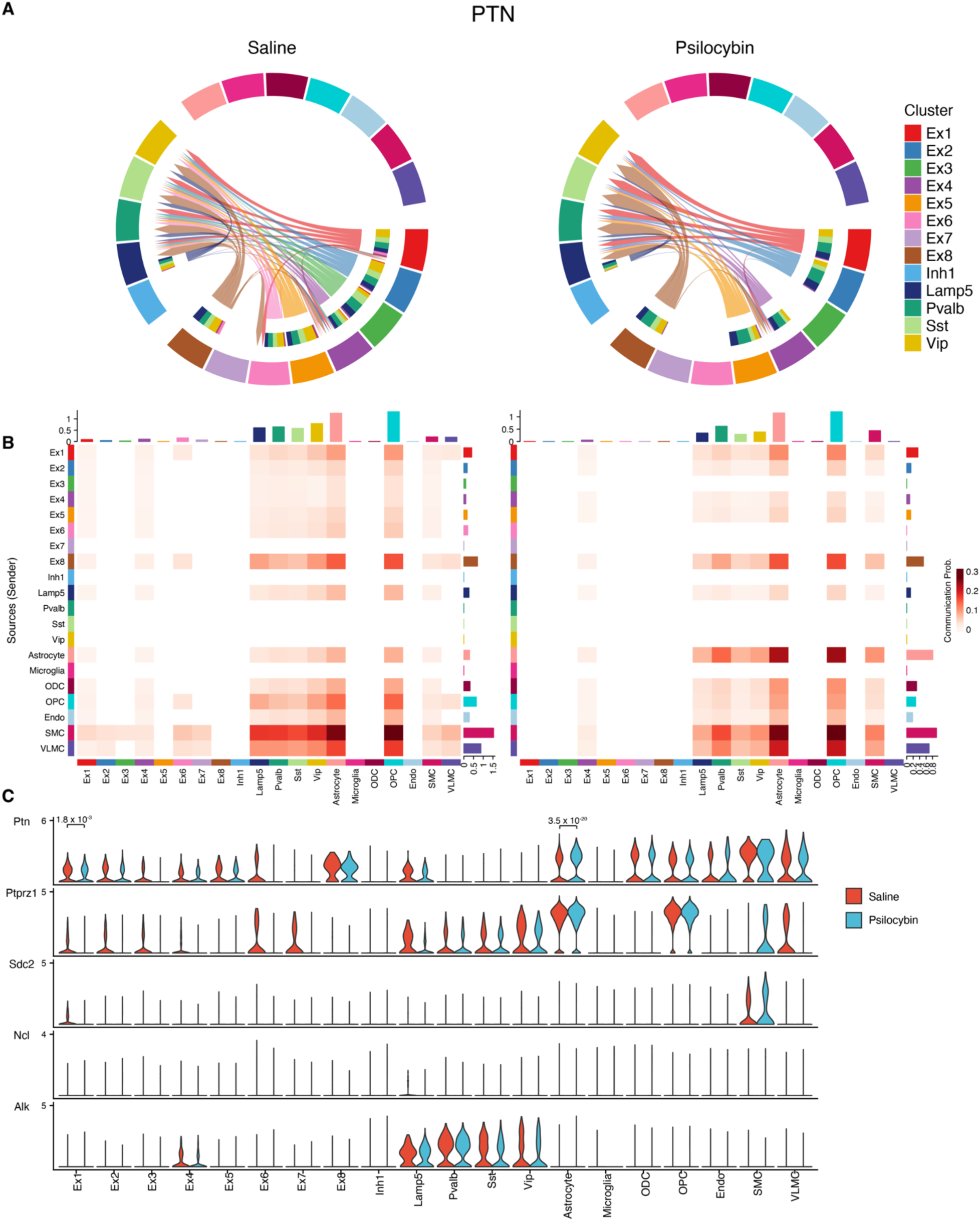
The comparison of the PTN signaling pathway of clusters between saline and psilocybin mice, related to Figure 2. **(A)** Chord diagrams of the PTN signaling pathway interaction strength of neuron clusters of saline and psilocybin mice. **(B)** The heatmaps of the PTN signaling pathway communication probability between cell clusters of saline and psilocybin mice. **(C)** The comparison of the PTN signaling pathway-related gene expression distribution in clusters of saline and psilocybin mice. *P*, Wald test with Benjamini-Hochberg procedure, calculated with DESeq2.

**Figure S15.**
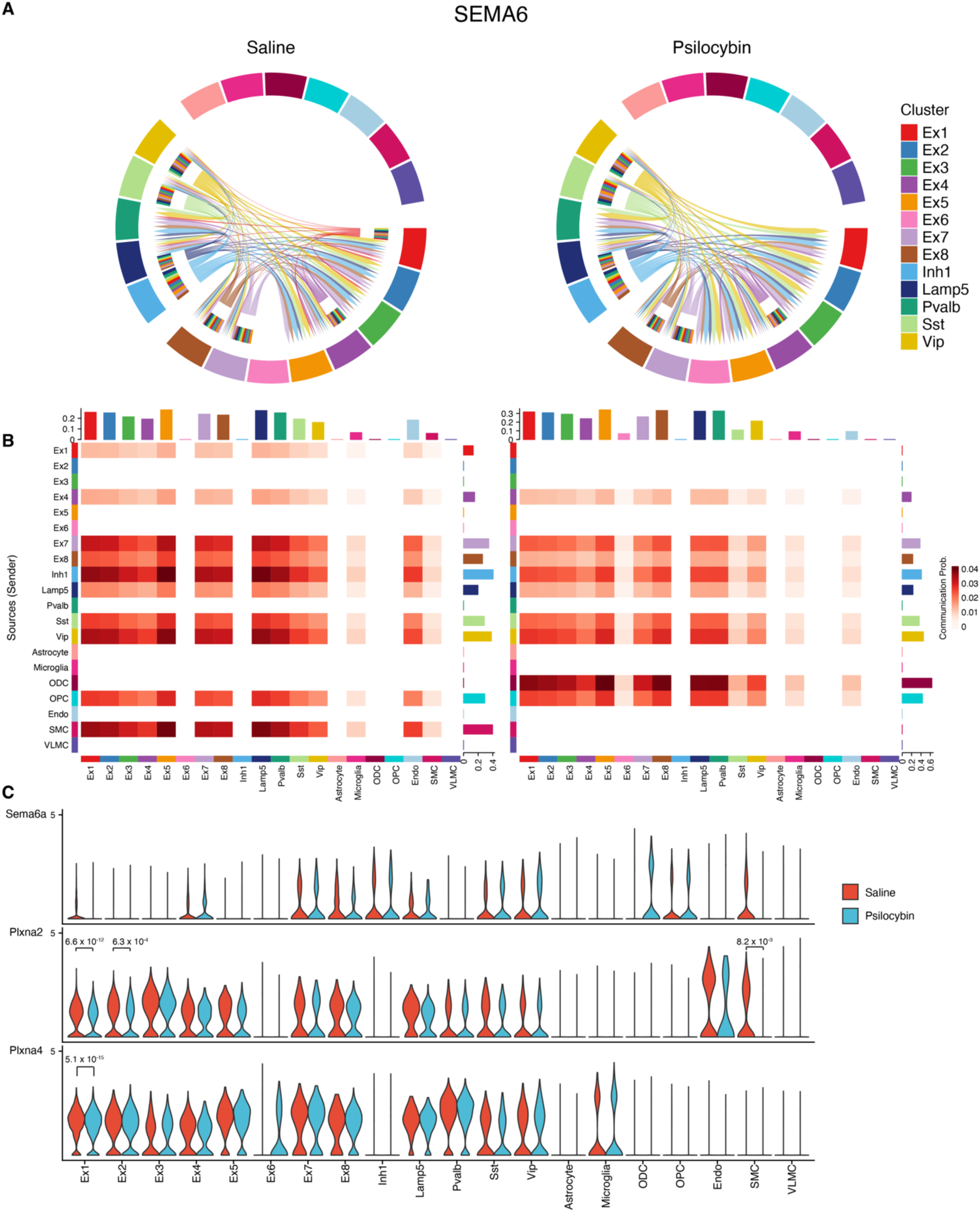
The comparison of the SEMA6 signaling pathway of clusters between saline and psilocybin mice, related to Figure 2. **(A)** Chord diagrams of the SEMA6 signaling pathway interaction strength of neuron clusters of saline and psilocybin mice. **(B)** The heatmaps of the SEMA6 signaling pathway communication probability between cell clusters of saline and psilocybin mice. **(C)** The comparison of the SEMA6 signaling pathway-related gene expression distribution in clusters of saline and psilocybin mice. *P*, Wald test with Benjamini-Hochberg procedure, calculated with DESeq2.

**Figure S16.**
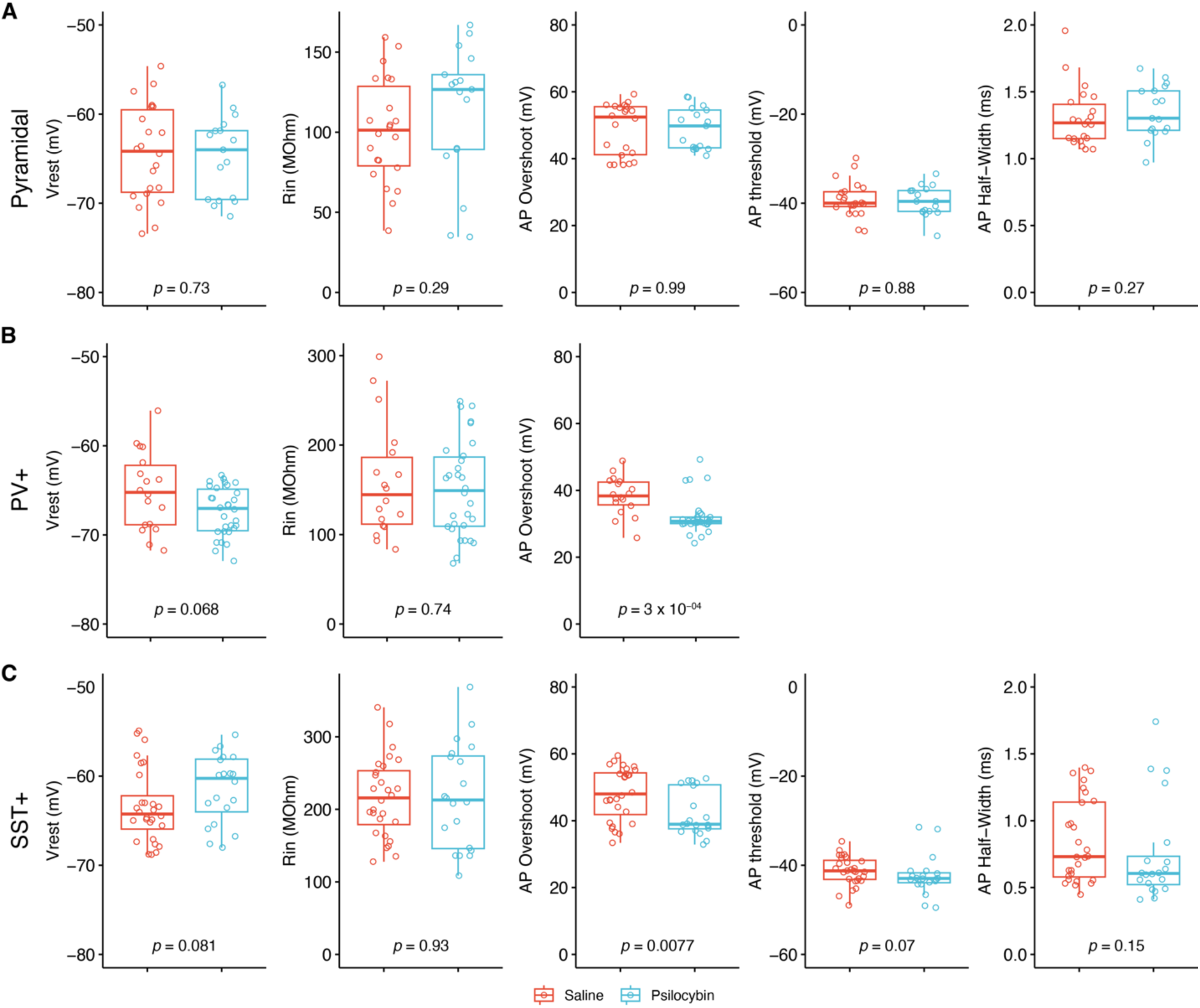
Statistical analyses of the intrinsic properties of layer 5 pyramidal, PV+, and SST+ neurons, related to Figure 3. **(A)** Comparisons of the resting membrane potential (Vrest), input resistance (Rin), overshoot of action potential (AP Overshoot), threshold of action potential (AP threshold), and half-width of action potential (AP Half-Width) of layer 5 pyramidal neurons in the OFC between Saline and Psilocybin mice (*p*, Wilcoxon test; Saline, n = 22, N = 4; Psilocybin, n = 17; N = 6). **(B)** Comparisons of the resting membrane potential (Vrest), input resistance (Rin), and overshoot of action potential (AP Overshoot) of PV+ neurons in the OFC between Saline and Psilocybin mice (*p*, Wilcoxon test; Saline, n = 18, N = 5; Psilocybin, n = 30; N = 7). **(C)** Comparisons of the resting membrane potential (Vrest), input resistance (Rin), and overshoot of action potential (AP Overshoot) of Sst+ neurons in the OFC between Saline and Psilocybin mice (*p*, Wilcoxon test; Saline, n = 28, N = 3; Psilocybin, n = 20; N = 3).

**Figure S17.**
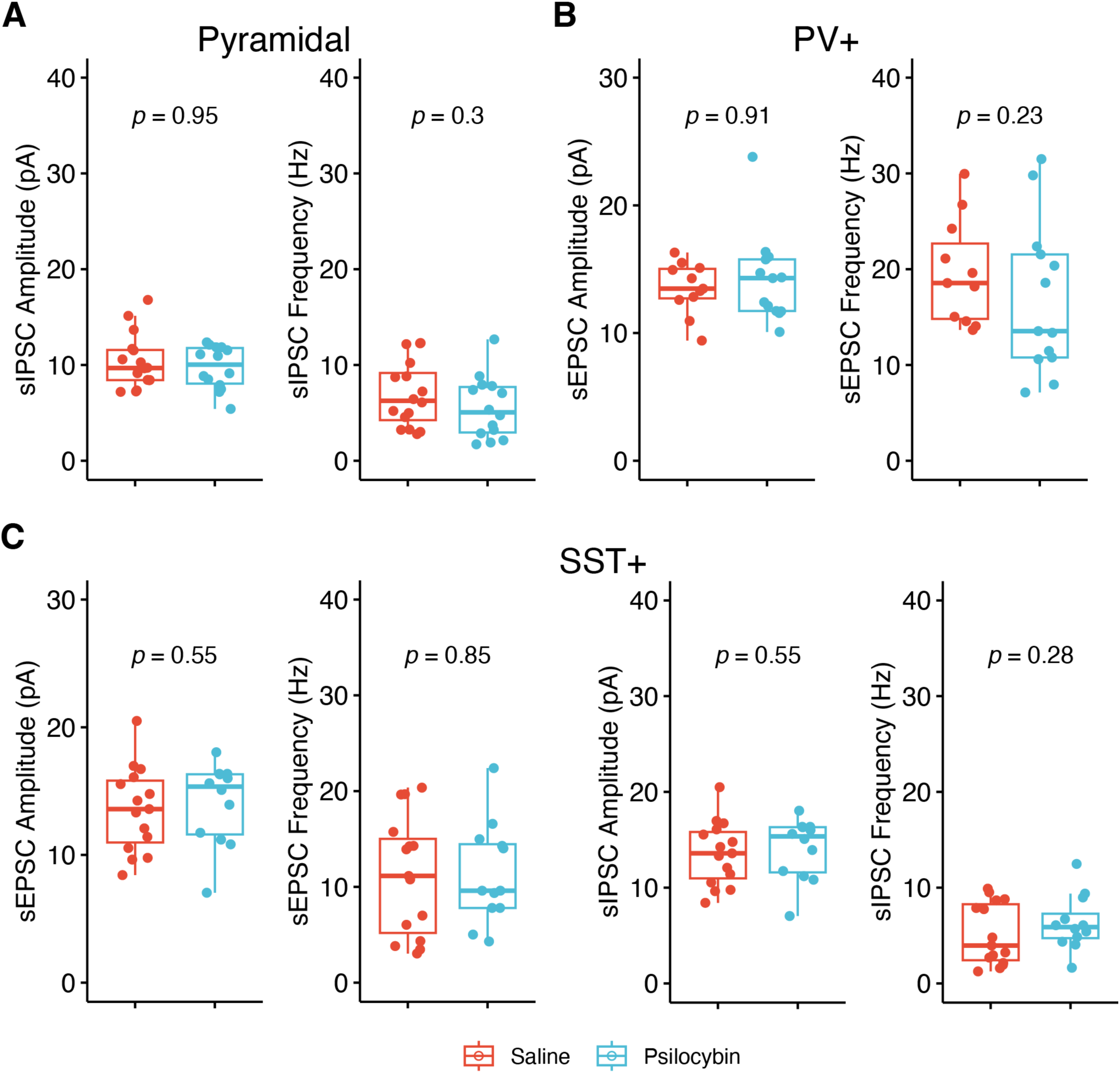
Statistical analyses of synaptic transmissions of the layer 5 pyramidal, PV+, and SST+ neurons, related to Figure 3. **(A)** Comparisons of the spontaneous inhibitory synaptic transmission of layer 5 pyramidal neurons in the OFC of Saline and Psilocybin mice (*p*, Wilcoxon test; Saline, n = 22, N = 4; Psilocybin, n = 20; N = 5) **(B)** Comparisons of the spontaneous inhibitory synaptic transmission of PV+ neurons in the OFC of Saline and Psilocybin mice (*p*, Wilcoxon test; Saline, n = 14, N = 6; Psilocybin, n = 12; N = 6). **(C)** Comparisons of the spontaneous excitatory and inhibitory synaptic transmission of PV+ neurons in the OFC of Saline and Psilocybin mice (*p*, Wilcoxon test; Saline, n = 22, N = 3; Psilocybin, n = 23; N = 3).

**Figure S18.**
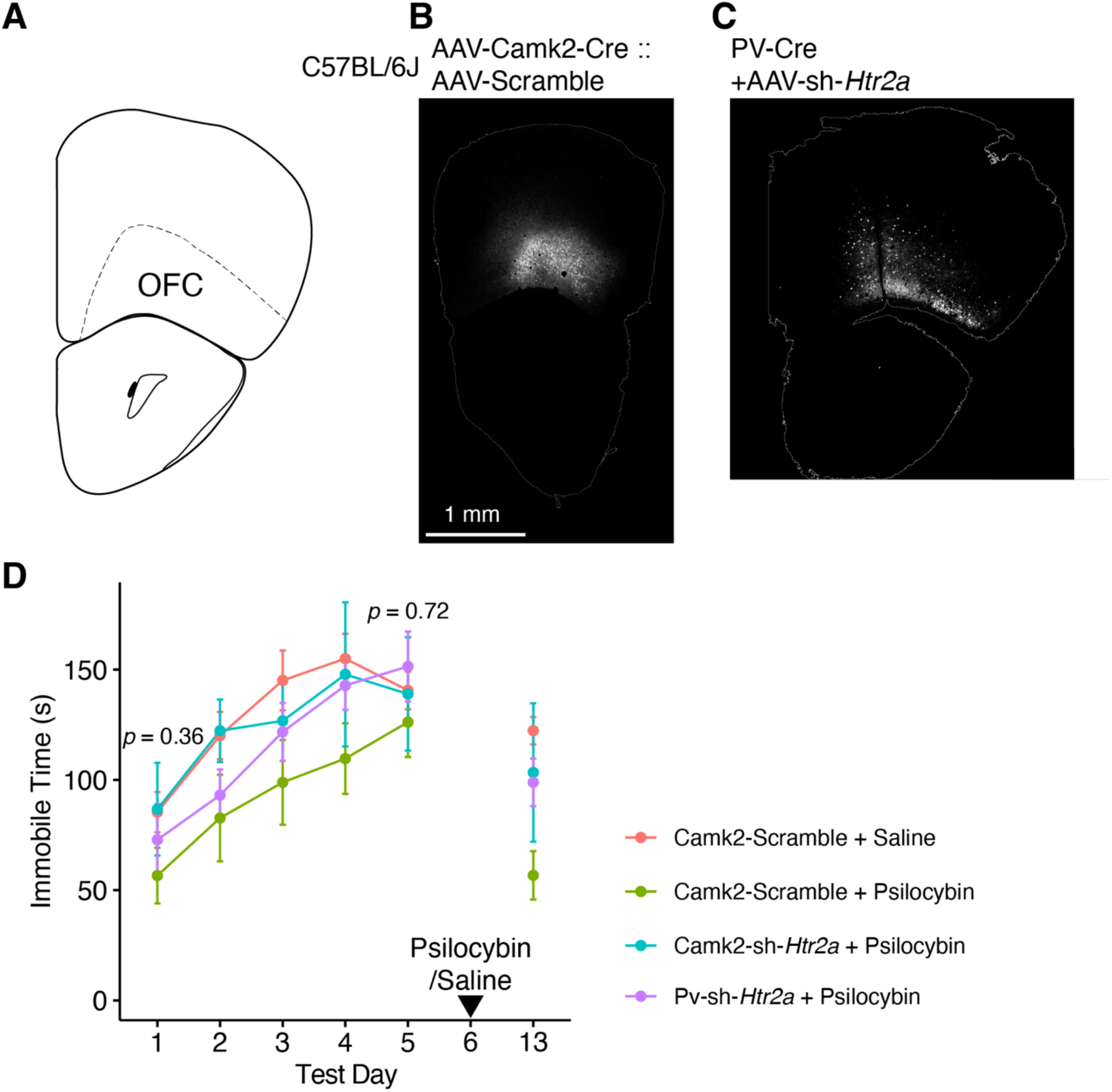
Knockdown of *Htr2a* blocked anti-depressant effect of psilocybin in rFST, related to Figure 4. **(A -C)** Representative viral infection in the OFC 2 weeks after viruses injections. **(D)** The time course of immobile time in FSTs, related to **Figure 4C**. Psilocybin or saline was *<u>i.p.</u>* injected 24 hrs after the FST on day 5 (Test Day 1: One-way ANOVA, F_(3,29)_ = 1.11, *p* = 0.36; Test Day 5: One-way ANOVA, F_(3,29)_ = 0.44, *p* = 0.72; Saline, N = 10; Psilocybin, N = 10; Camk2-sh-*Htr2a*, N= 6; PV-sh-*Htr2a*, N = 7).

**Table S1.** Summary of Cell Distribution in Clusters, related to Figure 1.

| **Cluster Name** | **Cell Number** | **Percentage of Total Cells (%)** | **Saline** | **Psilocybin** |
| --- | --- | --- | --- | --- |
| **Glutamatergic** |  |  |  |  |
| Ex1 | 3,681 | 14.59 | 1,347 | 2,334 |
| Ex2 | 2,224 | 8.81 | 809 | 1,415 |
| Ex3 | 924 | 3.66 | 392 | 532 |
| Ex4 | 785 | 3.11 | 412 | 373 |
| Ex5 | 745 | 2.95 | 316 | 429 |
| Ex6 | 519 | 2.06 | 354 | 165 |
| Ex7 | 365 | 1.45 | 136 | 229 |
| Ex8 | 362 | 1.43 | 114 | 248 |
| **GABAergic** |  |  |  |  |
| Pvalb | 1,040 | 4.12 | 461 | 579 |
| Vip | 966 | 3.83 | 444 | 522 |
| Sst | 731 | 2.90 | 320 | 411 |
| Inh1 | 831 | 3.29 | 308 | 523 |
| Lamp5 | 383 | 1.52 | 136 | 247 |
| **Non Neuronal** |  |  |  |  |
| Astrocyte | 4,676 | 18.53 | 2,785 | 1,891 |
| Endo | 619 | 2.45 | 561 | 58 |
| SMC | 486 | 1.93 | 411 | 75 |
| VLMC | 669 | 2.65 | 340 | 329 |
| OPC | 2,030 | 8.04 | 1,034 | 996 |
| Microglia | 1,528 | 6.05 | 806 | 722 |
| Oligodendrocytes | 1,673 | 6.63 | 760 | 913 |
| **Total** | **25,237** | **100.00** | **12,246** | **12,991** |

**Table S2.**
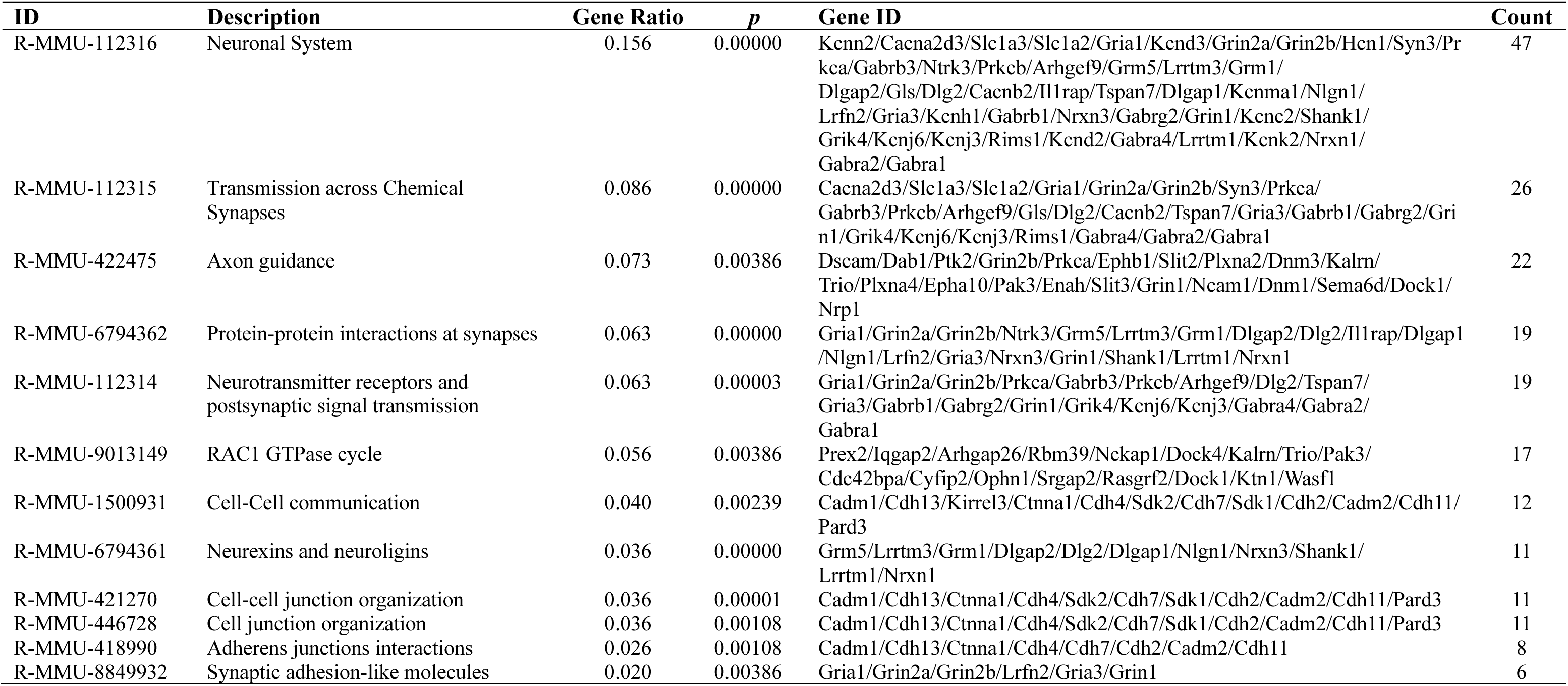
Reactome pathway-based gene enrichment analysis, related to Figure 1.

## Methods

### Mice

C57BL/6J mice (RRID:IMSR_JAX:000664) and PV-Cre driver strain (B6.Cg-*Pvalb^tm1.1(cre)Aibs^*/J, RRID:IMSR_JAX:012358) were used. Mice genotyping was performed following the guidance of the Jackson Laboratory. Adult male mice aged 7 – 10 weeks-old were used, except for PV-Cre mice in repeated forced swimming test, PV-Cre mice aged 12 – 16 weeks-old were used. Mice were maintained on a 12-hour light/dark cycle with food and water *ad libitum*. All experiments were performed in the dark cycle.

All procedures are in accordance with the National Institutes of Health *Guide for the Care and Use of Laboratory Animals*, and have been approved by Peking University Animal Care and Use Committee.

### Single-nucleus RNA Sequencing (snRNA-seq) analysis

Mice were anesthetized with isoflurane and decapitated, the orbitofrontal cortex was dissected on ice, then froze and stored at -80 ℃ before further experiments. Due to the relative low number of nucleus extracted from the sample of an individual mouse, samples of each experiment group were pooled together (N = 4/group). Nuclei were isolated following 10x Genomics single cell protocols, suspended and loaded into the 10x Genomics controller. Single cell GEMs and libraries generation were performed according to Chromium Next GEM Single Cell 3ʹ kit workflow, then sequenced on the 10x Genomics sequencing platform with Illumina NovaSeq 6000.

#### Sequence Alignment and UMI counting

Cellranger (v7.0.1, RRID:SCR_023221) analysis pipeline was used to process raw reads. Briefly, Cellranger mkfastq was used to convert raw base call (BCL) files of the snRNA-seq libraries into FASTQ files with a mouse Reference genome (mm10) as per the instructions provided by 10x Genomics. Cellranger was used for alignment, filtering, barcode counting, and unique molecular identifier counting.

#### snRNA-seq Analysis

Filtered count matrices from each sample were analyzed with Seurat (v4.3.0, RRID:SCR_016341) ^67^ and scCustomize (v1.1.3) ^68^ in R (v4.2.2, RRID: SCR_001905).

To remove low quality cells and likely multiplet captures, we filtered out cells with features < 700, mitochondrial gene percentage > 5%, then removed doublets with DoubletFinder (v2.0.3, RRID:SCR_018771) ^69^. Possible estimation and removal of cell free mRNA contamination was removed with SoupX (v.1.6.0, RRID:SCR_019193) ^70^. Gene expression differences were identified between Saline and Psilocybin samples (N = 4/group) using DESeq2 (v1.38.0, RRID:SCR_015687) ^71^. The enrichment analysis was performed with clusterProfiler (v4.8.1, RRID:SCR_016884) ^72^ and ReactomePA (v1.44.0, RRID:SCR_019316) ^73^.

For cell-cell interaction analysis, CellChat (v1.6.0, RRID:SCR_021946) was used to infer, visualize, and analyze intercellular communications from snRNA-seq data ^39^. CellChat basically utilize a signaling molecule interaction database of the known structural composition of ligand-receptor interactions, such as multimeric ligand-receptor complexes, soluble agonists and antagonists, as well as stimulatory and inhibitory membrane-bound co-receptors. With it CellChat infers cell-state specific signaling communications within scRNA-seq datasets using mass action models, along with differential expression analysis and statistical tests on cell groups, quantitatively characterize and compare the inferred intercellular communications through social network analysis tool, pattern recognition methods and manifold learning approaches.

### Virus injection and stereotaxic surgery

Adeno-associated viruses (AAV) were used in the present study. They were AAV2/9-SST-mCherry-WPRE-bGHpolyA (titer: 5.6 x 10^12^ GC/mL, BrainVTA), AAV/9-CaMKIIα-Cre (titer: 2.0 x 10^12^ GC/mL, BrainCase), and AAV/9-EF1α-DIO-mCherry (titer: titer: >=5.0 x 10^12^ GC/mL, BrainCase), AAV-CaMKIIα-EYFP-WPRE-PA(titer: 5.33 x 10^12^ GC/mL, BrainVTA), AAV/9-EF1α-DIO-EGFP-5’miR-30a-shRHA(mHtr2a)-3’miR-30a (titer: >=5.0 x 10^12^ GC/mL, BrainCase), and AAV/9-EF1α-DIO-EGFP-5’miR-30a-shRHA(Scramble)-3’miR-30a (titer: >=5.0 x 10^12^ GC/mL, BrainCase). The sequences of sh-Htr2a and Scramble are GCTCTGTGCCGTCTGGATTTA and CCTAAGGTTAAGTCGCCCTCG, respectively.

AAV viruses were injected into the orbitofrontal cortex of mice. Mice were anesthetized with 2 % isoflurane in oxygen at a flow rate of 2 L/min and mounted on a stereotaxic frame (RWD). Mouse body temperature was maintained with a temperature control blanket at 37 °C. Sterile ocular lubricant ointment was applied to the corneas to prevent drying. The scalp fur was shaved, and the skin was cleaned with 70 % alcohol and betadine. A hole was drilled at the injection site (A/P: + 2.4 mm; M/L: 1.2 mm, DV: -2.5 mm) using a 0.5-mm diameter round burr on a high-speed rotary micromotor (RWD). A total of 250 nL of virus solution was injected at a rate of 50 nL/min using a micro pump and Micro4 controller (World Precision Instruments).

After the injection, the needle was kept in the parenchyma for 5 - 10 min before being slowly withdrawn. The hole was sealed with bone wax and the skin was sutured. Mice were placed in a 37 °C isothermal chamber to recover from anesthesia before returned to their home cage.

### RNAscope *in situ* hybridization assay and immunostaining

To analyze the expression of *Htr2a* mRNA neurons of the OFC, we performed RNA *in situ* hybridization to quantify mRNAs as described previously ^16^. Probes against the mRNAs of *Htr2a* (Advanced Cell Diagnostics, Cat#401291), *Slc17a7* (Cat#416631), *Sst* (Cat# 404631), and *Pvalb* (Cat# *421931*) were used. In brief, mouse brains were obtained and quickly frozen in isopentane on dry ice for 20 s maximum, and kept in -80 ℃ overnight or until use. Coronal brain sections of 16 μm thickness were collected in a cryostat at -20 °C. We next fixed those sections with 10 % neutral buffered formalin solution (Sigma, Cat#HT5011) for 20 min, followed by washing steps with PBS and dehydrating steps with alcohol. With the RNAScope H_2_O_2_ and Protease Reagents (Advanced Cell Diagnostics, Cat#322381), we treated sections with H_2_O_2_ for 10 min before with protease IV for 30 min in room temperature. After washing off the protease, we incubated brain sections with a mix of probe sets targeting mRNAs mentioned above for 2 hr at 40 °C in the HybEZ oven (Advanced Cell Diagnostics, RRID:SCR_012481). Following probe incubation, with the RNAscope Multiplex Fluorescent Assay v2 (Advanced Cell Diagnostics, Cat#323100) and RNAScope 4-Plex Ancillary kit for Multiplex Fluor (Advanced Cell Diagnostics, Cat#323120), sections went through a series of incubations with preamplifier probes, amplifier probes, and TSA Vivid Fluorophore 520 (Advanced Cell Diagnostics, Cat#PG-323271), TSA Vivid Fluorophore 570 (Advanced Cell Diagnostics, Cat#PG-323272), or TSA Vivid Fluorophore 650 (Advanced Cell Diagnostics, Cat#PG-323273) at 40 °C, and then counter-stained with DAPI.

For immunostaining, mice were anesthetized with isoflurane, and then perfused with phosphate-buffer saline (PBS, pH 7.4) and then 4% paraformaldehyde (PFA) in PBS. Brain tissues were post-fixed with 4% PFA in PBS overnight at 4 °C, and 25-μm coronal sections were prepared with a vibratome. Immunostaining followed the standard protocols for free-floating sections. In brief, free-floating sections were incubated in blocking solution containing 4% normal donkey serum, 1% bovine serum albumin (BSA), and 0.3% Triton X-100 in PBS, with slow shaking for 2 h at 23–25 °C. Sections were then treated with primary antibody in blocking solution for 24 – 48 hr at 4 °C and with secondary antibody in blocking solution at 23–25 °C for 2 h with slow shaking.

Primary antibodies used were Rabbit Anti-HTR2A (1:400, Thermo Fisher, Cat# PA5-120747, RRID:AB_2914319), Rabbit Anti-Glutamate receptor 1 (1:150, Millipore, Cat# AB1504, RRID:AB_2113602), Rat Anti-Somatostatin (1:200, Millipore, Cat# MAB354, RRID:AB_2255365), Goat Anti-Parvalbumin (1:2000, Swant, Cat# PVG-213, RRID:AB_2650496), Goat Anti-Iba1 (1:100, Abcam, Cat# ab289874, RRID:AB_2942069), Mouse Anti-Glial Fibrillary Acidic Protein (1:100, Millipore, Cat# MAB360, RRID:AB_11212597), Mouse Anti-NeuN (1:1000, Abcam, Cat# ab104224, RRID:AB_10711040), and Rat Anti-NeuN (1:2000, Abcam, Cat# ab279297).

Secondary antibodies used were Alexa Fluor Plus 647 Anti-Rabbit (1:300, Thermo Fisher, Cat# A32795, RRID: AB_2762835), Alexa Fluor 546 Anti-Goat(1:300, Thermo Fisher, Cat# A11056, RRID: AB_2534103), Alexa Fluor 594 Anti-Rabbit (1:300, Jackson ImmunoResearch Labs, Cat# 711-585-152, RRID: AB_2340621), Alexa Fluor Plus 647 Anti-Rat (1:300, Thermo Fisher, Cat# A48272, RRID: AB_2893138), Alexa Fluor 488 Anti-Mouse (1:300, Thermo Fisher, Cat# A21202, RRID: AB_141607), Alexa Fluor 405 Anti-Mouse (1:300, Abcam, Cat# ab175658, RRID: AB_2687445), and Alexa Fluor 405 Anti-Rat (1:300, Thermo Fisher, Cat# A48268, RRID: AB_2890549).

We acquired fluorescent images with a confocal microscope (Leica TCS-SP8 STED) using a 63× objective (NA 1.4). The maximal projection of a 5 μm thick stack was analyzed with ImageJ (v1.53t, RRID:SCR_003070) based FIJI (RRID:SCR_002285) ^27^. The analysis was performed as previously described ^56^. For *in situ* hybridization assay, the combined region of *Slc17a7, Pvalb,* or *Sst* and enclosed DAPI area was defined as the cell area. To count the density of the puncta, fluorescent dot numbers were measured and normalized to the cell area.

### Electrophysiological Recordings

To label PV+ and SST+ positive neurons, we injected AAV-EF1α-DIO-mCherry or AAV-SST-mCherry into the OFC of *PV-Cre*+ or C57BL/6J mice, respectively, mice were allowed for at least 12 days to recover. Mice were *i.p.* injected with saline or psilocybin (1 mg/mL) randomly, and we performed electrophysiology experiments 24 hrs after the injection.

Recordings were performed as previously described ^56^. Briefly, adult male mice aged 9 – 12 wks-old were anesthetized with isoflurane and decapitated. Brains were removed and sectioned in cold (0 - 4 °C) cutting solution (in mM): 87 NaCl, 3.0 KCl, 1.5 CaCl_2_, 1.3 MgCl_2_, 1.0 NaH_2_PO_4_, 26 NaHCO_3_, 20 D-glucose, and 75 sucrose, saturated with 95 % O_2_ and 5 % CO_2_ to obtain 250 μm-thick coronal sections with a vibratome (Leica VT1200S). Slices were transferred and incubated in a holding chamber containing (ACSF, in mM): 124 NaCl, 3.0 KCl, 1.5 CaCl_2_, 1.3 MgCl_2_, 1.0 NaH_2_PO_4_, 26 NaHCO_3_, and 20 D-glucose, saturated with 95% O_2_ and 5% CO_2_ at 33 °C for 30 min and then at room temperature for at least 30 min before recordings.

Layer 5 pyramidal neuron in the orbitofrontal cortex (OFC) was identified by cell morphology and size, and recorded using methods as previously described (Zhang et al., 2017), Parvalbumin- and Somatostatin-positive neurons were identified with fluorescence under the microscope (Olympus BX51WI). Brain slices were placed in a submersion type chamber continuously perfused with ACSF saturated with 95% O_2_ and 5% CO_2_ at 31 – 33 °C.

For spontaneous synaptic transmission recordings, the pipette solution contained the following (in mM): 120 Cs gluconate, 4 ATP-Mg, 0.3 GTP, 0.5 EGTA, 10 HEPES, and 4.0 QX-314 (pH 7.2, 270–280 mOsm with sucrose).

For current-clamp recording, the pipette solution contained (in mM): 120 potassium gluconate, 10 KCl, 4 ATP-Mg, 0.3 GTP, 10 HEPES, and 0.5 EGTA (pH 7.2, 270–280 mOsm with sucrose). Series resistance was fully compensated using the bridge circuit of the amplifier MultiClamp 700B (Molecular Devices, RRID: SCR_018455). Action potential threshold was estimated as the point when the slope of rising membrane potential exceeds 50 mV ms^−1^.

Electrodes had resistances of 2 ∼ 3.5 MΩ. The series resistance was not compensated in voltage-clamp experiments. During experiments, the series resistance was constantly monitored. Recordings were discarded when series resistance was >16 MΩ or change of series resistance was >15%. Recordings were made with Multiclamp 700B amplifier and PCIe-6321 (National Instruments, RRID:SCR_018947) controlled by AxoGraph X (AxoGraph). Data were filtered at 4 kHz and digitized at 20 kHz. Data were analyzed offline with Axograph X and NeuroMatic package ^74^ (RRID: SCR_004186) in Igor Pro (Wavemetrics, RRID: SCR_000325).

### Behavior tests

#### Repeated forced swimming test

The repeated forced swimming test was conducted as described in previous studies ^52–54^. Briefly, the mice were forced to swim for 5 min daily for five consecutive days and then tested again on day 13. On day 6 and 24 hours after the fifth forced swimming test, the mice were *i.p.* injected with 0.2 ∼ 0.3 ml saline or 1 mg/kg body weight psilocybin dissolved in saline. Mice were gently transferred into a glass beaker (18 cm diameter, 13 cm water height at 26 ± 2 °C water). The behavior was recorded and analyzed offline by an observer blind to the treatment. After test, animals were dried with towel and placed in a warmer cage till fur fully dried, then returned to home cages.

#### Open field test and head-twitch Response

Prior to testing, mice were transported to the test room and left undisturbed for at least 30 minutes. Animal activity in a plastic test arena (40 x 40 x 40 cm) was recorded with a camera mounted above the center. Mice were placed in the center of the arena for open field test. The activities of mice over 10 mins were recorded. For open field test the first 5 mins of animal activities in the arena were analyzed with Matlab (MathWorks, RRID:SCR_001622). The central area was defined as the 30 x 30 cm square space in the middle of the arena. The variables measured were distance travelled, time in the perimeter versus in the central area. For head twitches, the video clips of the first 10 mins of animal activities in the arena were scored.

### Statistical analysis

All reported sample numbers represent biological replicates. All statistical analyses and data plotting were performed with R (v4.2.2, RRID: SCR_001905), and the non-base attached packages for R were ggpubr (v0.6.0, RRID:SCR_021139), rstatix (v0.7.2, RRID:SCR_021240), tidyverse (v2.0.0, RRID:SCR_019186), emmeans (v1.8.5, RRID:SCR_018734). For boxplots, whiskers denoted 1.5 * IQR from the hinges, which corresponded to the first and third quartiles of distribution. For comparisons between two groups, non-parametric Wilcoxon test was used. For comparisons between multiple groups, Shapiro-Wilk test was used for normality test, and Levene’s test was used for variance test. For multiple groups, two-way ANOVA with *post-hoc* Tukey’s test or one-way ANOVA with *post hoc* Holm-Bonferroni procedure were used based on experiment design. n, sample number of cells; N, sample number of mice. *P* < 0.05 is considered statistically significant.

### Drug

Psilocybin (Cayman #14041) was dissolved in sterile saline and administered at a dose of 1.0 mg/mL intraperitoneally.

### Data Availability

The raw and processed snRNA-seq data reported in this paper are available online (GEO: GSE246451). Any additional information that supports the findings of this study are available from the lead contact upon request.

